# Flexible cue anchoring strategies enable stable head direction coding in both sighted and blind animals

**DOI:** 10.1101/2022.01.12.476111

**Authors:** Kadjita Asumbisa, Adrien Peyrache, Stuart Trenholm

## Abstract

Vision plays a crucial role in instructing the brain’s spatial navigation systems. However, little is known about how vision loss affects the neuronal encoding of spatial information. Here, recording from head direction (HD) cells in the anterior dorsal nucleus of the thalamus in mice, we find stable and robust HD tuning in blind animals. In contrast, placing sighted animals in darkness significantly impairs HD cell tuning. We find that blind mice use olfactory cues to maintain stable HD tuning and that prior visual experience leads to refined HD cell tuning in blind adult mice compared to congenitally blind animals. Finally, in the absence of both visual and olfactory cues, the HD attractor network remains intact but the preferred firing direction of HD cells continuously drifts over time. We thus demonstrate remarkable flexibility in how the brain uses diverse sensory information to generate a stable directional representation of space.

**Highlights:**

- Head direction (HD) cell tuning in ADn is robust in blind animals, but unstable in sighted animals placed in the dark
- Blind mice use olfaction to stabilize HD cell tuning
- Prior visual experience leads to refined HD cell tuning in blind adult mice
- In the absence of both vision and olfaction, the HD attractor network in ADn remains intact but the preferred firing direction of HD cells continuously drifts

## Introduction

Our visual system provides critical and up-to-date information about the world around us, facilitating navigation through the environment and enabling quick reactions to dynamic events. Vision facilitates navigation related tasks including landmarking, obstacle avoidance and the generation of an internal cognitive spatial map (Tolman, 1948; O’Keefe, 1979; Klatzky, 1998; Schinazi et al., 2016). In the absence of vision, other sensory modalities need to fill the previously dominant role of vision in generating spatial awareness and guiding navigation. In humans, while it is clear that visually-impaired individuals can successfully form spatial maps and navigate in many environments, behavioral studies have noted differences in various aspects of spatial awareness and navigation between sighted and visually-impaired individuals (Thinus-Blanc and Gaunet, 1997; Pasqualotto and Proulx, 2012; Schinazi et al., 2016). However, little is known about how the brain’s spatial navigation systems, which have been best examined in freely moving rodent studies (Moser et al., 2017), adapt following vision loss.

The brain has dedicated systems for generating spatial awareness and guiding spatial navigation. In rodents, some of the most studied components of the brain’s spatial awareness system include place cells (O’Keefe, 1979), grid cells (Hafting et al., 2005) and head direction cells (Taube et al., 1990a) – though other complementary spatially-tuned cells also exist (Moser et al., 2017). To examine the effect of vision loss on the brain’s spatial navigation system, we wanted to focus on a brain region from which we could reliably record from a high percentage of spatially-coding cells in order to maximize our chances of measuring possible similarities/differences in spatial tuning between sighted and blind animals. We thus decided to focus on head direction (HD) cells in the anterior dorsal nucleus (ADn) of the thalamus, where the spike rate of a majority of neurons is modulated by head direction (Taube, 1995; Peyrache et al., 2015).

The head direction system encompasses a network of interconnected brain regions that incorporate angular head velocity signals with additional sensory information about environmental landmarks in order to generate cells with highly tuned and stable preferences for heading direction (Taube, 2007). Ablation/silencing studies have shown that intact HD cells are required for proper spatial navigation (Wilton et al., 2001; Butler et al., 2017; Harvey et al., 2017) and accurate spatial representation within the brain’s other spatial navigation systems (Winter et al., 2015; Harland et al., 2017). The HD system appears to be organized as an attractor network (Skaggs et al., 1995; Peyrache et al., 2015; Chaudhuri et al., 2019; Angelaki and Laurens, 2020), such that pairs of simultaneously recorded HD cells maintain a similar angular difference between their preferred firing directions across exposures to different rooms and following environmental manipulations that alter tuning preferences (Taube et al., 1990b). As such, cells with similar HD tuning preferences tend to fire coherently even when tuning preferences are unstable (Bassett et al., 2018) or during sleep (Peyrache et al., 2015; Chaudhuri et al., 2019).

Although vestibular inputs are critical for the HD signal (Stackman and Taube, 1997; Stackman et al., 2002; Muir et al., 2009), vision also appears to play an important role in anchoring and providing stability to the HD system. For example, horizontally displacing a visual cue in an environment reliably causes a concomitant shift in the preferred firing direction (PFD) of HD cells (Taube et al., 1990b). However, many studies have found that HD cell responses are relatively stable in the absence of visual inputs or visual cues (Blair and Sharp, 1996; Chen et al., 2016; Butler et al., 2017; Dannenberg et al., 2020). Furthermore, recordings from HD cells in young rodents just before eye opening revealed tuned HD cells, though tuning curves were found to be broader than following eye opening (Bjerknes et al., 2015; Tan et al., 2015; Bassett et al., 2018).

It has thus been argued that the HD system only requires idiothetic inputs (i.e. internally generated sensory signals, such as vestibular, proprioceptive, or motor efference copy signals), with vision only providing a refining allothetic (i.e. externally generated) sensory input (Goodridge et al., 1998; Stackman et al., 2003; Taube, 2007). In contrast, other studies have found highly unstable HD cell responses following removal of visual inputs in adult mice (Mizumori and Williams, 1993; Yoder and Taube, 2009), arguing for the possible requirement of external sensory inputs in stabilizing HD tuning. Additionally, while it is clear that vision can exert a strong effect on HD cell tuning preferences, the extent to which other sensory systems can provide landmarking cues and stabilize HD cell tuning remains unclear. While rodents have been found capable of using both auditory and olfactory cues to guide spatial learning and navigation (Watanabe and Yoshida, 2007; Fischler-Ruiz et al., 2021; Poo et al., 2021), previous studies with HD cells have suggested that auditory, olfactory, and vibrissal systems are relatively ineffective in modulating HD cell tuning (Goodridge et al., 1998; Long and Zhang, 2021). Thus, it remains uncertain what effect vision loss might have on HD cell responses. Here, to examine the effect of vision loss on HD cells in adult mice, we recorded from HD cells in ADn of both sighted and blind animals, and explored the extent to which HD cell responses were altered following vision loss and whether other sensory systems could be leveraged to tune the HD system.

## Results

### Robust head direction tuning in blind mice

Is HD cell tuning affected by vision loss? To examine this, we recorded from neurons in ADn of rd1 mice (Chang et al., 2002), a rodent model of retinitis pigmentosa in which mice are born with normal vision but go completely blind by ∼ 1 month old, due to photoreceptor degeneration (Stasheff et al., 2011). For all experiments, recording probes were implanted in ADn of adult mice (2-4 months old) and animals were placed in a circular open field arena (Figure 1A; 60 cm diameter; black walls with a single visual cue; see Methods). The animal’s position and heading direction were tracked with a set of IR video cameras while spiking responses of HD cells were recorded (see Methods). Following experiments, spike sorting was performed to isolate individual HD cells, based on methods previously described for identifying HD cells in sighted animals (see Methods). Unless otherwise noted, HD cell responses were analyzed over the entirety of 10-minute-long recording sessions (see Methods).

**Figure 1.**
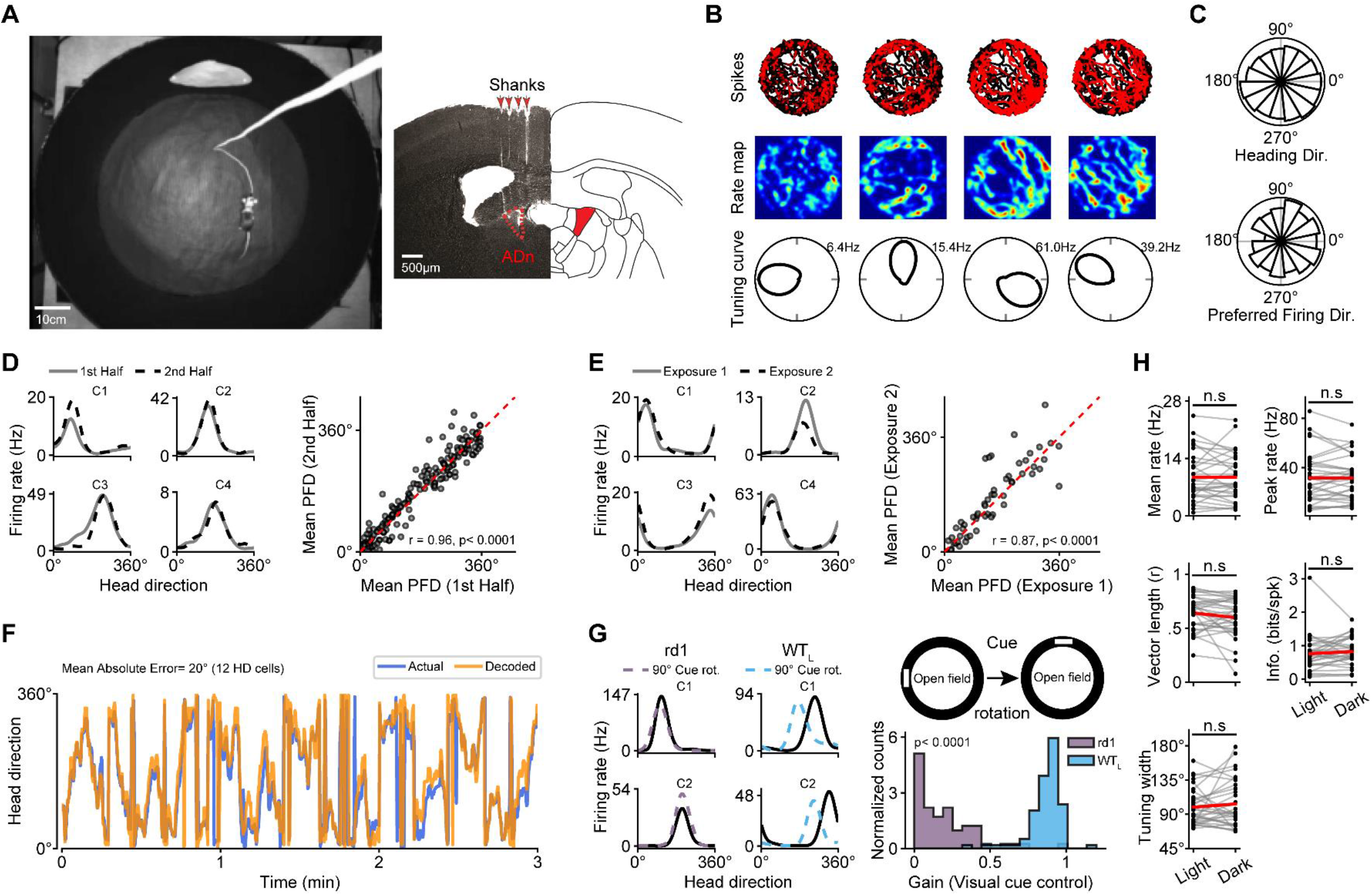
Robust and stable head direction cells in ADn of blind mice. **A**, *left*, A picture of the open field recording arena, featuring a mouse with an electrode implant. The arena was 60cm in diameter, with black walls and a single visual cue. **A**, *right*, An example post hoc coronal brain slice showing the tracts of the 4-shank recording electrode (*left*), with the anterior dorsal nucleus (ADn) of the thalamus indicated in red, and a corresponding slice of the brain atlas shown (*right*). **B**, *top*, The position of an rd1 mouse in the open field arena over a 10-minute recording session is shown in black, and the spatial locations that evoked spiking responses are shown for 4 simultaneously recorded HD cells from an rd1 mouse are indicated in red. **B***, middle*, For the same cells as above, heat maps in which the spike rate is normalized according to position occupancy, showing that the recorded cells do not have preferred spatial firing locations. **B**, *bottom*, For the same 4 cells as above, polar plots indicating the preferred firing directions. **C**, *top*, A polar plot showing the occupancy of angular bins for heading direction of all rd1 mice over a 10-minute recording session (see Methods). **C**, *bottom*, A polar showing the preferred firing directions (PFDs) for HD cells recorded across all rd1 mice (see Methods). **D**, *left*, The spike rate vs. heading direction of four simultaneously recorded HD cells (C1-C4) in an rd1 mouse during the first and last 5 minutes of a 10-minute recording session. **D**, *right*, For all HD cells across animals (151 HD cells across 13 animals), the mean PFD is compared between the first and last 5 minutes. The data points closely follow the diagonal (red dotted line). The pre-post tuning similarity was tested with a Pearson correlation, and the resulting *r* value is shown. **E**, The same as in **D**, except comparing HD cell tuning across successive 10-minute exposures to the same open field arena (55 HD cells across 7 animals). **F**, An analysis comparing the actual heading direction of an rd1 mouse over time (blue) to the heading direction predicted by a Bayesian decoder (orange, see Methods). **G**, The spike rate vs. heading direction of 2 simultaneously recorded HD cells in an rd1 (*left*) and sighted (*middle*) mouse in control conditions (solid black line) and following a 90° visual cue rotation (dotted line). **G**, *right, A* schematic indicating the nature of visual cue rotation experiments, and a histogram showing the extent of control (gain) that visual cue rotation exerted on the PFD of HD cells in rd1 vs. sighted mice (see Methods; statistical difference was tested with the Mann-Whitney U Test). WT_L_ = 93 HD cells across 6 animals; rd1 = 65 HD cells across 6 animals. **H**, Several metrics characterizing HD cells in rd1 mice are compared during light vs. dark exposure. Statistical differences were calculated using the Wilcoxon Signed-Rank Test. n.s = not statistically different. rd1 = 31 HD cells across 4 animals in both light and dark conditions.

Upon placing blind adult rd1 mice in the open field environment, we found robust HD cell tuning (Figure 1B; n = 151 HD cells recorded from 13 animals, with 80.7 % of recorded cells in ADn defined as HD cells; see Methods). Similar to what has been described for sighted animals, we found that rd1 mice evenly sampled all angular directions in the environment and possessed HD cells that exhibited preferred firing directions that spanned all heading directions (Figure 1C).

We next tested the stability of HD cells in blind mice, and the extent to which their HD cell network was able to reliably encode heading direction. First, we compared the preferred direction of HD cells in the first and second half of 10-minute recording sessions, and found that preferred firing directions remained stable over a single session (Figure 1D). Second, we examined the stability of preferred firing directions across repeated exposure to the same room (see Methods), and found that preferred direction tuning was highly stable across repeated exposure (Figure 1E). These stability metrics were similar to those measured in sighted animals in the light (Supplementary Figure 1). Next, we examined whether the HD cell network in ADn of blind mice was providing a reliable readout of the animal’s heading direction by performing a decoding analysis (Zhang et al., 1998; Johnson et al., 2005). We found robust and stable decoding of HD with as few as 12 simultaneously recorded HD cells (Figure 1F). Therefore, in blind mice HD cells are highly tuned and provide an accurate readout of heading direction.

To ensure that rd1 animals were indeed blind, we performed visual cue rotation experiments in the light. Upon rotating a visual cue, HD cells in rd1 mice failed to follow the cue, unlike HD cells from sighted animals in the light (Figure 1G). Furthermore, various metrics that we computed for HD cells in rd1 mice (e.g. mean and peak firing rates, resultant vector length and tuning width) were statistically similar in both light and dark environments (Figure 1H). Thus, despite a total absence of rod and cone based visual inputs, HD cell tuning is robust and stable in blind animals, providing accurate information about heading direction, meaning that blind mice can generate a stable allocentric spatially-guided map.

### HD cell tuning is more robust in blind animals than in sighted animals placed in the dark

How do HD cell responses in blind animals compare to those of sighted animals? To test this, we recorded HD cell responses from sighted animals in the light (WT_L_) and in complete darkness (WT_D_; see Methods; Figure 2A). A similar percentage of cells recorded in ADn passed the criteria to be designated as HD cells in rd1 and WT_L_ mice (Figure 2B; see Methods), whereas a significantly smaller percentage of ADn cells were designated as HD cells in WT_D_ mice (Figure 2B). Thus, HD cell tuning is significantly more robust is blind animals than sighted animals placed in the dark, meaning that blind animals have adapted an alternative non-visual strategy for stabilizing HD cell tuning.

**Figure 2.**
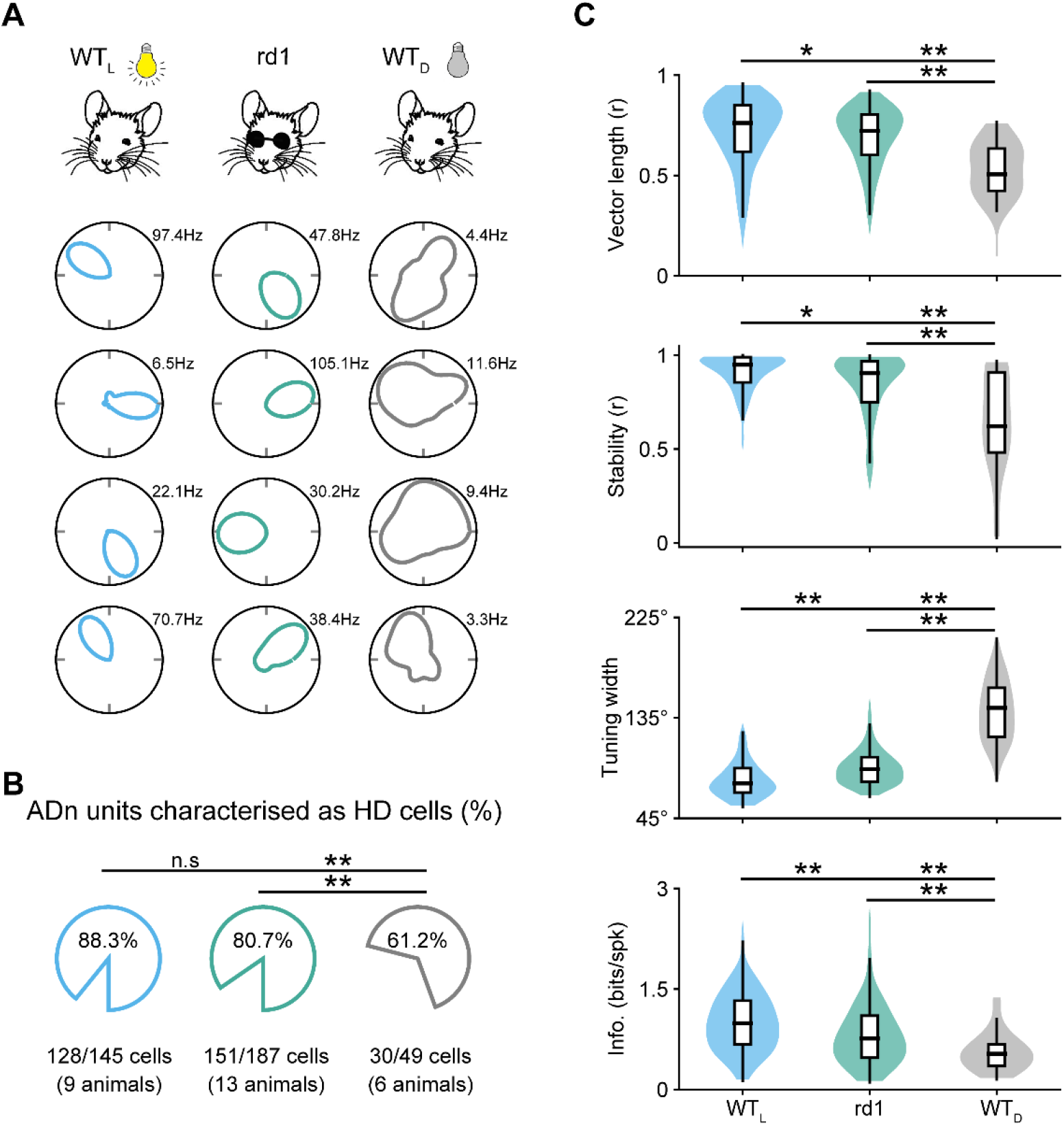
HD cell tuning is more robust in blind animals than in sighted animals placed in the dark. **A**, For 3 different groups of mice – wildtype in light (WT_L_; blue), rd1 (blind; green), and wildtype in the dark (WT_D_; grey) – polar plots of 4 simultaneously recorded HD cells are shown. **B**, For the 3 different groups of mice outlined in A, the average percent of cells recorded in ADn that passed the criterion to be designated as HD cells (see Methods) are compared. Statistical differences were calculated using the Z-test for Proportions. **C**, For the 3 different groups of mice outlined in **A** and **B**, several characteristics of HD cells are compared. For each graph, data are shown as hybrid violin/box plots (see Methods). Additional statistical comparisons are provided in Supplementary Figure 2. Statistical differences were calculated using the Kolmagorov-Smirnov Test with Bonferroni Correction for multiple comparisons.

Next, we compared several metrics for HD cell responses between blind and sighted mice: mean and peak firing rates were similar between rd1, WT_L_ and WT_D_ mice (Supplementary Figure 2). In contrast, for vector length, tuning width, stability and mutual information, each group of mice was statistically different from one another, demonstrating the following hierarchy in HD tuning refinement: WT_L_ > rd1 > WT_D_ (Figure 2C). Thus, sighted mice exhibit significant impairment in HD cell tuning when placed in the dark, with a level of impairment much more severe than is seen in blind animals, for whom HD cell tuning – though slightly less refined than in sighted animals in the light – is robust and stable.

### Prior visual experience leads to refined HD cell tuning in blind adult mice

We next tested whether vision might be required during development for proper maturation of HD cell tuning, even if mice subsequently go blind. For instance, in the superior colliculus, it has been shown that normal vision is required during development for proper formation of auditory maps (King et al., 1988; King and Carlile, 1993). In mice, eye opening occurs ∼ P10-12 (Gordon and Stryker, 1996; Smith and Trachtenberg, 2007), and rd1 mice have normal vision for a few days following eye opening, followed by some amount of attenuated vision for approximately 2 more weeks until they go fully blind around 1 month old (Stasheff et al., 2011). To test whether vision around the time of eye opening is required to refine and mature the tuning of the HD system, we performed experiments in Gnat2^cpfl3^ Gnat1^irdr^/Boc mice (subsequently referred to as Gnat1/2^mut^), who are congenitally blind due to dysfunctional rod and cone photoreceptors (see Methods). Recording from adult Gnat1/2^mut^ mice (2-4 months old), we found many highly tuned HD cells (Figure 3A; n = 128 HD cells recorded from 8 animals) with a similar percentage of cells in ADn designated as HD cells for Gnat1/2^mut^ mice as we found for both rd1 and WT_L_ mice, but with Gnat1/2^mut^ mice exhibiting a significantly higher percentage of HD cells than WT_D_ mice (Figure 3B; Supplementary Figure 3). Similar to rd1 mice, HD cells in Gnat1/2^mut^ mice maintained stable preferred directions throughout a 10-minute session, as well as across repeated exposure to the same environment (Supplementary Figure 3). Furthermore, similar to rd1 mice, HD cells in Gnat1/2^mut^ mice failed to follow the visual cue when the cue was moved. Under both light and dark conditions, their HD tuning did not exhibit any noticeable change (Supplementary Figure 3). Finally, at the population level, simultaneously recorded HD cells provided a reliable decoding of the animal’s heading direction (Supplementary Figure 3). However, based on the metrics that we calculated from HD cell responses during exploration of the open field environment, we found that HD cell tuning was statistically less refined in Gnat1/2^mut^ mice compared to rd1 mice on several metrics tested (Figure 3C; Supplementary Figure 2). Nonetheless, HD cell tuning in Gnat1/2^mut^ mice was significantly more refined than in sighted animals placed in the dark (WT_D_; Supplementary Figure 2), indicating the following hierarchy in HD cell tuning refinement: WT_L_ > rd1 > Gnat1/2^mut^ > WT_D_. These results indicate that while visual inputs are not required for generating stable HD tuning in adult blind mice – either in rd1 or Gnat1/2^mut^ animals – normal visual inputs in the days after eye opening appear to be required for refinement and maturation of vision-independent HD tuning in adult blind mice.

**Figure 3.**
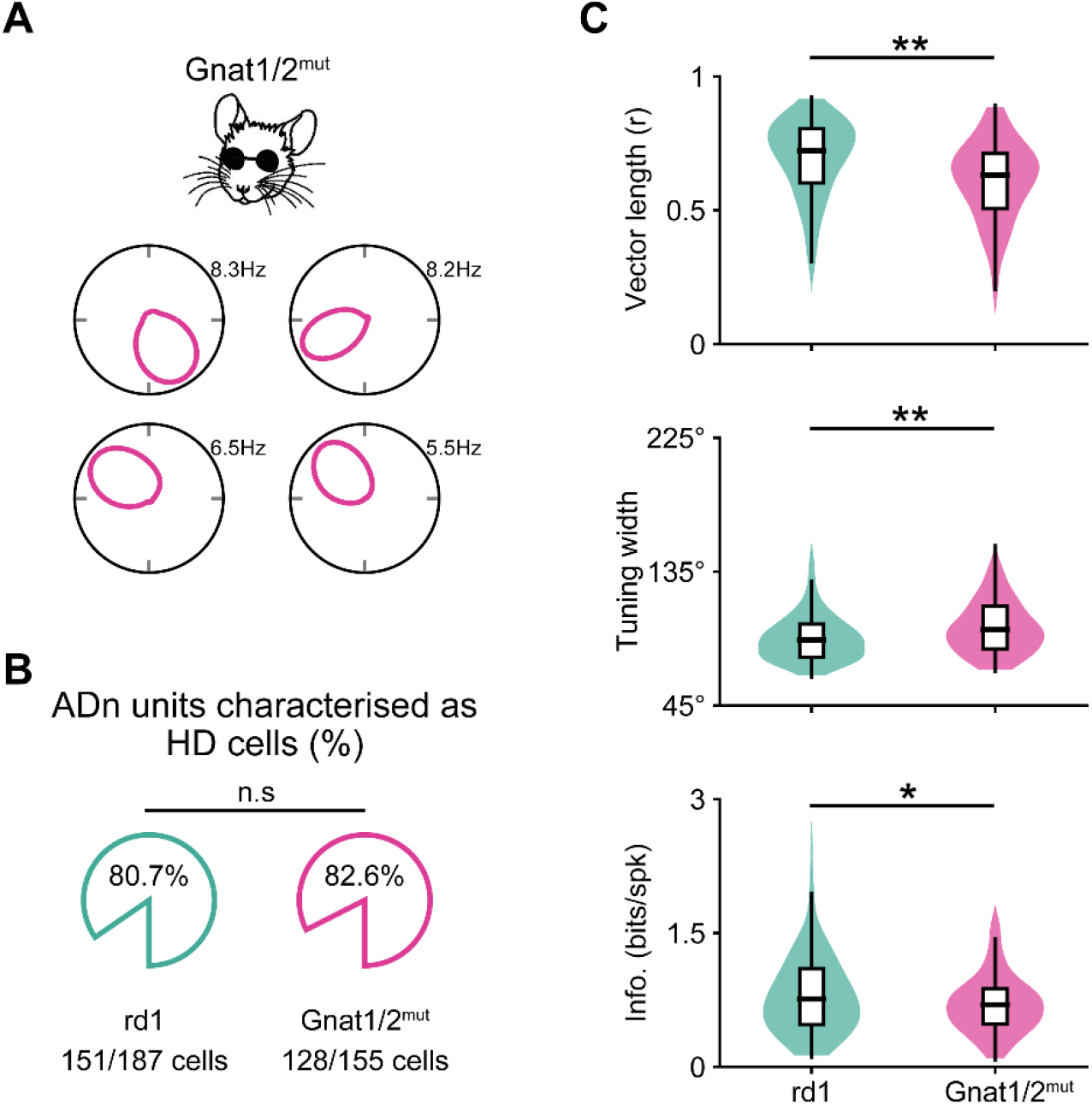
Prior visual experience leads to refinement of HD cell tuning in blind adult mice. **A**, For a Gnat1/2^mut^ mouse (congenitally blind; pink), polar plots of 4 simultaneously recorded HD cells are shown. **B**, For rd1 (replotted from Figure 2) and Gnat1/2^mut^ mice, the average percent of cells recorded in ADn that passed the criterion to be designated as HD cells (see Methods) are compared. Statistical differences calculated using the Z-test for Proportions. For Gnat1/2^mut^ mice, 128 HD cells across 8 animals. **C**, Several characteristics of HD cells are compared between rd1 (replotted from Figure 2) and Gnat1/2^mut^ mice. For each graph, data are shown as hybrid violin/box plots (see Methods). Statistical differences were calculated using the Kolmagorov-Smirnov Test. Additional statistical comparisons are provided in Supplementary Figure 2.

### The vibrissal system is not required for head direction cell tuning in blind animals

We next tested whether blind mice were using an alternative sensory modality to landmark their HD system within the open field environment. As all our experiments were performed with a speaker playing white noise placed immediately underneath the center of the open field arena, we deemed it unlikely that auditory inputs were contributing to HD cell tuning in our experiments (see also Goodridge et al., 1998). We first examined the vibrissal system, hypothesizing that a mouse might be surveying the walls and floor of the open field arena with its whiskers to aid in spatial awareness. To test this, we shaved the whiskers from blind mice (see Methods) and examined the effect on HD cell responses (Figure 4A). Whisker shaving did not result in any noticeable change in HD cell tuning, except for a small reduction in the peak firing rate (Figure 4A; we pooled together rd1 and Gnat1/2^mut^ mice as HD cells in both were similarly unaffected by whisker shaving (Supplementary Figure 4)). Thus, the vibrissal system does not appear to play a major role in HD cell tuning in blind animals.

**Figure 4.**
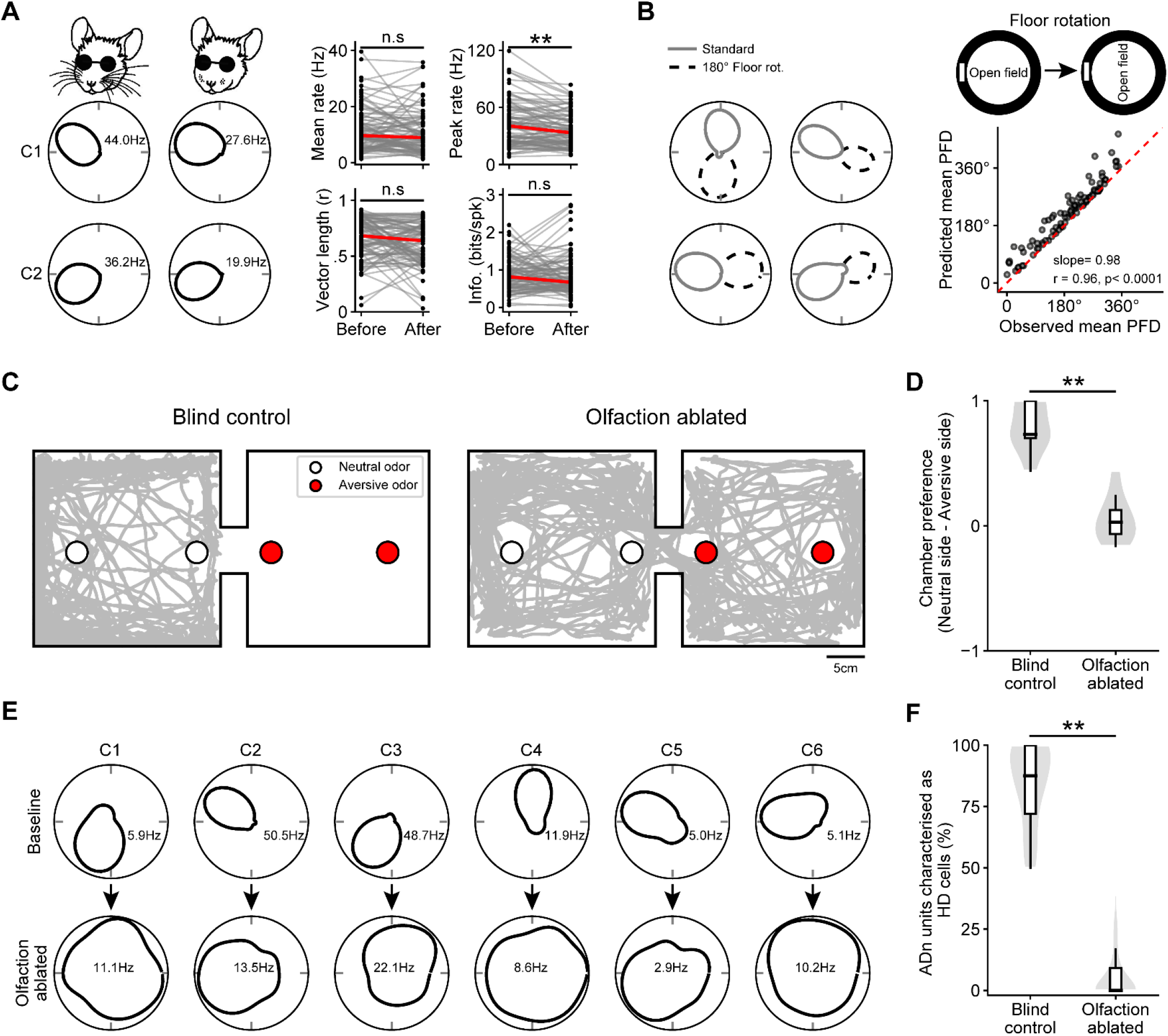
Olfactory signals are required for stable HD cell tuning in blind animals. **A**, Example polar plots of 2 simultaneously recorded HD cells in an rd1 mouse before (*left*) and after (*middle*) whisker ablation. **A**, *right*, Several HD cell metrics are compared before and after whisker ablation, pooled for rd1 and Gnat1/2^mut^ (independent analyses for these different mouse lines are shown in Supplementary Figure 4). Statistical comparisons were computed using the Wilcoxon Signed Rank Test. Blind = 89 HD cells across 11 animals. **B**, *left*, Polar plots from 4 simultaneously recorded HD cells in an rd1 mouse before (solid gray line) and after (dotted black line) the floor was rotated by 180° (see Methods). **B**, *right (top)*, Schematic of the floor rotation experiment. **B**, *right (bottom)*, Upon floor rotation, the observed shift in the mean PFDs is plotted against the expected shift in mean PFDs (see Methods; pooled for rd1 and Gnat1/2^mut^ (independent analysis for these different mouse lines is shown in Supplementary Figure 4)). The r value listed was computed with the Pearson correlation on 94 HD cells across 9 animals. **C**, Example of an olfactory test, in which an rd1 mouse was placed in a 2-room chamber, with one room containing an aversive odor (3-methyl-1-butanethiol) and the other room containing a neutral odor (distilled H_2_0). The trajectory of the mouse over a 10-minute recording session is plotted in grey in control (*left*) and following olfactory sensory neuron ablation (*right*). **D**, The time spent in the neutral room is compared to time spent in the aversive room across blind animals. Statistical test used was the Mann Whitney U Test. Blind control = 8 animals; Olfaction ablated = 9 animals. **E**, Polar plots for 6 simultaneously recorded HD cells in an rd1 mouse that were recorded in control condition (*top*) and the next day following olfactory sensory neuron ablation (*bottom*; see Methods). **F**, Violin/box plots are shown comparing the percent of recorded cells in ADn that passed the criterion for being designated as HD cells (see Methods) in control and following olfactory ablation. Statistical test used was the Mann Whitney U Test.

### The olfactory system is required for stabilizing head direction cell tuning in blind animals

Due to the stability of preferred firing directions observed across repeated exposures that we had previously noted for blind mice, we suspected they might be using olfaction to anchor the HD system. To test this, we performed a floor rotation experiment. Here, instead of rotating a visual cue on the wall as we did previously for visual cue rotation experiments, we rotated the floor (i.e. the animal was removed from the room, the floor was rotated without being cleaned, and the animal was reintroduced to the room). In blind mice, floor rotation resulted in a concomitant shift in the preferred firing direction of HD cells (Figure 4B; we pooled together rd1 and Gnat1/2^mut^ mice as results were similar between these two blindness models (Supplementary Figure 4)), suggesting that olfaction could modulate HD cell tuning. In blind animals, floor rotation led to concomitant shifts in the preferred direction of HD cells at levels similar to that observed in sighted animals in light following visual cue rotations (Supplementary Figure 4).

To directly test whether olfactory inputs were required for stabilizing HD cell tuning in blind mice, we ablated olfactory sensory neurons (OSNs) and examined the effect on HD cell tuning. To ablate OSNs, we used a previously established chemical lesioning method (Norwood et al., 2019; see Methods). To ensure that this method was effective at ablating OSNs and severely impairing olfaction, we developed an olfactory place avoidance task in which a mouse was placed in a two-chamber arena, with one of the chambers housing an aversive olfactory substance (3-methyl-1-butanethiol (Sievert and Laska, 2016)) and the other chamber housing a neutral olfactory substance (distilled H_2_O; Figure 4C; see Methods). Mice with intact OSNs completely avoided the chamber containing the aversive olfactory substance (Figure 4C,D). In contrast, following OSN ablation, mice showed no preference between the two chambers (Figure 4C,D). Having established a method to reliably ablate olfaction, we tested the effect of olfactory ablation on HD cell tuning in blind animals. Olfactory ablation resulted in complete loss of HD cell tuning in blind mice (Figure 4E,F; we pooled together rd1 and Gnat1/2^mut^ mice as OSN ablation affected both similarly (Supplementary Figure 4)). The percent of ADn cells that passed the criteria for being designated as HD cells drastically decreased for blind mice following olfactory ablation (Figure 4F). For some experiments, whiskers were intact during olfactory ablation, while in other animals whiskers were ablated prior to olfactory ablation, but in both cases olfactory ablation resulted in complete loss of HD cell tuning (Supplementary Figure 4), again indicating that the vibrissal system was not being used to tune HD cells. These results show that blind mice use olfactory cues to anchor their HD signal.

### Olfaction can modulate HD cell tuning in sighted animals

We next examined whether olfaction could also impact HD cell tuning in sighted animals. First, we noted that HD cell tuning was more robust in sighted animals in the dark (WT_D_) than in blind animals following OSN ablation (Figure 5A; Supplementary Figure 5; see Methods for a description of an alternate method we developed to define HD cells in blind mice following OSN ablation once HD cell tuning curves became unstable). We thus wondered whether the weak but remnant tuning of HD cells in sighted animals in the absence of visual inputs could be arising from olfactory cues. To test this, we performed floor rotation experiments. Similar to blind animals, HD cells in sighted animals in the dark consistently rotated towards the direction of floor rotation (Figure 5B; Supplementary Figure 5). However, floor remapping was significantly weaker in WT_D_ compared to blind mice, with HD cells in WT_D_ mice consistently and significantly undershooting the full extent of the floor rotation (Figure 5B; Supplementary Figure 5). Olfaction therefore has an impact on HD cell tuning in sighted animals, though it appears much less effective at tuning HD cell responses than in blind animals. To test this more directly, we ablated OSNs in sighted animals. Following olfactory ablation, the percentage of cells recorded in ADn that passed the criteria to be designated as HD cells significantly decreased in WT_D_ (Figure 5C) and HD cell responses became similar to those of blind animals following olfactory ablation (Supplementary Figure 5). In contrast, following olfactory ablation the percentage of recorded cells in ADn that passed the criteria to be designated as HD cells was unaffected in WT_L_ (Figure 5D; Supplementary Figure 5). Thus, sighted animals can use both vision and olfaction to provide cue anchoring information to the HD system, though at least in the open field environment we used for experiments vision appears to be more effective at tuning the HD system. Only in the absence of both visual and olfactory cues does the HD system become completely untuned.

**Figure 5.**
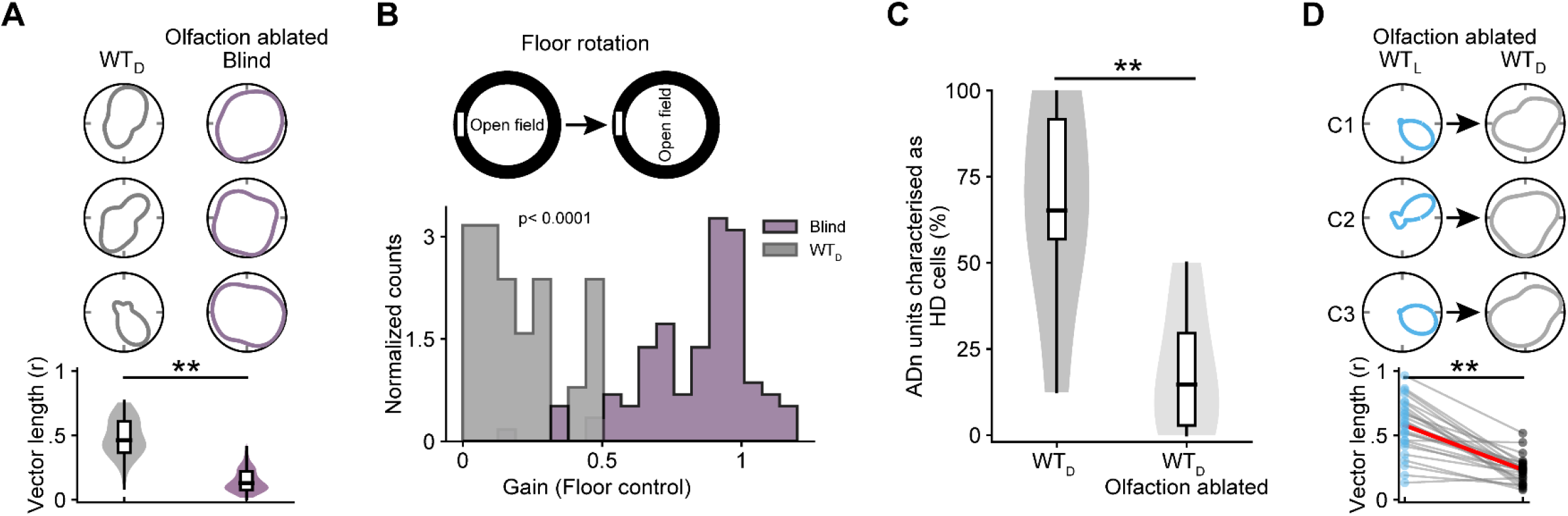
Olfaction modulates HD cell tuning in sighted mice placed in the dark. **A**, top, Example polar plots from 3 simultaneously recorded HD cells in a sighted animal placed in the dark (WT_D_, *left*) and a blind rd1 mouse following olfactory sensory neuron ablation (*right*). **A,** *bottom*, The vector length of tuning curves of HD cells belonging to blind animals following olfactory ablation is compared to that of sighted animals placed in the dark (42 HD cells for WT_D_; 106 HD cells for Blind; comparison of additional metrics is shown in Supplementary Figure 5). Statistical comparison computed with the Mann Whitney U Test. **B**, *top*, Schematic outlining the floor rotation manipulation. **B**, *bottom*, Histogram showing the extent of control the floor rotation exerted on the PFD of HD cells in sighted mice in the dark (WT_D_) vs. olfaction ablated blind mice (see Methods; blind pooled for rd1 and Gnat1/2^mut^ (independent analysis for these different mouse lines is shown in Supplementary Figure 5)). Statistical difference was tested with the Mann-Whitney U Test. WT_D_ = 93 HD cells across 5 animals. Blind = 94 HD cells across 9 animals. **C**, Violin/box plots are shown comparing the percent of recorded cells in ADn that passed the criterion for being designated as HD cells (see Methods) in sighted mice in the dark (WT_D_) before and after olfactory sensory neuron ablation. Statistical differences were calculated using the Z-test for Proportions. WT_D_ = 6 animals; WT_D_: Olfaction ablated = 5 animals. **D**, *top*, Example polar plots of 3 simultaneously recorded HD cells from a sighted animal following olfactory sensory neuron ablation, placed in either the light (*left*) or dark (*right*). **D**, *bottom*, Vector length of tuning curves for HD cells from sighted animals placed in both light vs. dark environments following olfactory sensory neuron ablation (comparison of additional metrics is shown in Supplementary Figure 5). 27 HD cells across 5 animals in both light and dark conditions.

### Attractor dynamics in the absence of allothetic (external) sensory cues

We next tested how HD attractor dynamics were affected following the loss of cue anchoring visual and olfactory signals. We found that in the absence of both visual and olfactory cues, while HD cells became completely untuned, the activity of simultaneously recorded cells in ADn were well-described by a one-dimensional ring manifold (Figure 6A), similar to the HD network in blind control animals (Figure 6A), and in sighted animals in the light (Supplementary Figure 6). However, following the loss of both vision and olfaction, the population activity on the ring manifold representing the moment-to-moment heading direction of the animal became decoupled from the actual heading direction (Figure 6A,B; Supplementary Figure 6). For such a ring manifold to persist in the absence of visual and olfactory inputs, the relative firing of HD cells with respect to one another must be maintained as the animal moves around the environment, even though each cell no longer maintains a stable preferred firing direction over time. How could this arise? Previous work has shown that prior to eye opening, while HD cell tuning is broad when measured over an extended period of time, if tuning curves are computed on short timescales then much more refined tuning curves can be measured (Bassett et al., 2018). Therefore, instead of computing tuning curves by averaging data across 10-minute recording sessions as we have done until now, following removal of both visual and olfactory inputs we computed tuning curves each time an animal made an angular head rotation of 360° (which occurred every 25.4 ± 10.3 seconds; see Methods). On these shorter intervals, we found that HD cells were highly tuned (Figure 6C,D) and similar to control conditions (Supplementary Figure 6), but that their preferred directions were constantly drifting over time, with all simultaneously recorded cells drifting in a coherent manner (Figure 6C). Thus, in the absence of cue anchoring sensory signals, though HD cell tuning becomes unstable over longer periods of time, the attractor network in ADn remains intact and HD cells exhibit sharp tuning curves over short time intervals.

**Figure 6.**
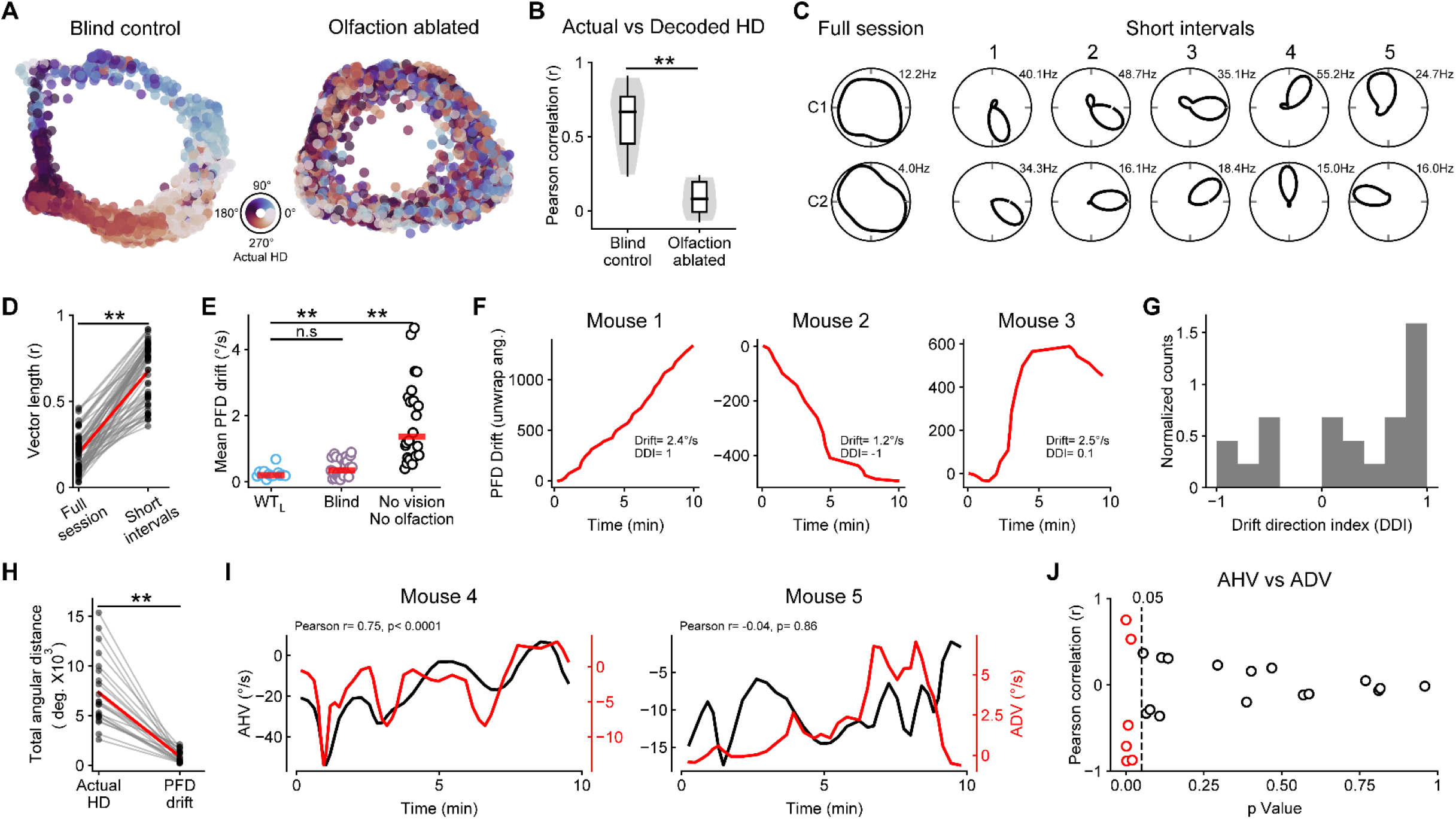
In the absence of vision and olfaction, the HD attractor in ADn remains intact but the ‘hill’ of activity drifts over time. **A**, Isomap plots representing the activity of the population of simultaneously recorded HD cells over time, for a control blind rd1 mouse (*left*) and the same animal following olfactory sensory neuron ablation (*right*). Each circle represents the low-dimensional location of the population activity at a given point in time. The location of points in this low dimensional space, plotted over time, form a 1-dimensional ring. Each dot is color coded based on the animal’s actual heading direction measured at that time point (additional isomap plots for the different types of mice and different manipulations used throughout this study can be found in Supplementary Figure 6). **B**, The relationship between a blind animal’s actual heading direction vs. the heading direction indicated by activity within the attractor is compared for blind mice before and after olfaction ablation. Statistical difference was tested with the Mann-Whitney U Test. **C**, Example polar plots for 2 simultaneously recorded HD cells in an rd1 mouse, either calculated over the entire 10-minute recording session (*left*) or over shorter timescales (*right*) with each successive epoch computed upon successive 360° head turns. **D**, The mean vector length computed from HD cell tuning curves is compared when measured for the entire 10-minute recording sessions or for the 360° head turn epochs. Statistical comparison with the Wilcoxon Signed-Rank Test. **E**, The average velocity of drift in the HD population is compared in sighted mice in the light (WT_L_), blind mice, and in mice without both visual and olfactory inputs. Statistical comparison with the Mann Whitney U Test. WT_L_ (median drift= 0.2°/s); Blind (median drift= 0.35°/s); No vision/olfaction (median drift= 1.36°/s). **F**, Example of HD cell population drift (i.e. the average PFD drift for all simultaneously recorded HD cells) from 3 example animals with no vision and olfaction, showing the rate and direction of PFD drift (positive values indicate CW drift; negative values indicate CCW drift). **G**, Histogram showing the extent that the overall drift direction was either CW (positive values) or CCW (negative values) for all mice with no vision and olfaction (see Methods, and Supplementary Figure 6 where the DDI calculated for different strains of mice is shown separately). **H**, The total angular distance covered for mice over the 10-minute recording session is compared to the total angular distanced covered by drift of the population of HD cell PFDs (for 14 animals). Statistical comparison with the Mann Whitney U Test. **I**, For 2 mice, in the absence of vision and olfaction, the angular head velocity (AHV) is compared to the angular drift velocity (ADV) of preferred firing directions. The Pearson correlation was computed between AHV and ADV curves. **J**, Scatter showing the Pearson r and p values computed for AHV versus ADV for all sessions where visual and olfactory inputs were blocked (as in panel **I**). Sessions where p < 0.05 and |r values| > 95^th^ percentile of shuffles are indicated in red (see Methods and Supplementary Figure 6).

Next, we quantified the drift of preferred firing direction of HD cells in the absence of visual and olfactory inputs. On average, we found that preferred firing directions within the HD network drifted between ∼1-4 degrees/s (see Methods), whereas in control conditions there was effectively no drift (Figure 6E; note that for the ‘No Vision and No Olfaction’ group we pooled together blind animals following olfactory ablation and sighted animals in the dark (WT_D_) following olfaction ablation (Supplementary Figure 6)). In some animals, the preferred firing directions of all simultaneously recorded HD cells drifted consistently in a clockwise (CW) direction over the 10-minute recording session (Figure 6F). In other animals, the preferred directions drifted consistently in a counter-clockwise (CCW) manner (Figure 6F), while for other animals the direction of drift alternated during the recording session (Figure 6F). To quantify the consistency in drift direction across a 10-minute recording session, we computed a drift direction index (DDI; see Methods) with a value of 1 indicating that the PFD of the HD population consistently drifted in a CW manner for the entire session and a value of −1 indicating that the PFD of the population consistently drifted in a CCW manner for the entire session. We found a large spread of DDI values, though there appeared to be a subtle bias for drifting in a CW direction (Figure 6G), particularly for blind animals (Supplementary Figure 6). Interestingly, within a given animal, bias in the direction of preferred direction drift appeared to be relatively stable across repeated exposures to the open field environment (Supplementary Figure 6).

Next, we examined if the drift in preferred firing direction of HD cells correlated with angular head movements. We found that over a 10-minute recording session, the animals rotated their heads significantly more than the PFD drift of the HD cell population (Figure 6H), indicating that the drift in PFD was not simply a one-to-one readout of head rotations. To better test if the animal’s head rotations were related to the direction and extent of PFD drift in the HD cell network, for each 360° head turn, we computed both the animal’s angular head velocity (AHV) and the preferred firing direction’s angular drift velocity (ADV; for each 360° head rotation we computed an HD cell tuning curve, and we then measured the angular distance between the preferred firing directions of subsequent 360° rotations, and divided by the time that transpired in between). We found that for some animals the AHV and ADV signals were significantly correlated, while for others there was no correlation (Figure 6I,J; Supplementary Figure 6). Therefore, while the HD attractor network is intact in the absence of visual and olfactory inputs, either visual or olfactory landmarking cues are required to anchor the attractor network and enable stable HD cell tuning. Without such allothetic external sensory cues, the PFD of HD cells is unanchored and constantly drifts. Finally, for some animals, the drift of the PFD appears to be coupled to ongoing angular head velocity signals, while for other animals it is unclear what is driving the drift.

## Discussion

Recording from head direction cells in ADn of both sighted and blind mice, we find stable and robust HD tuning in all animals. Remarkably, the HD system is flexible in which sensory system it can use for obtaining reliable cue anchoring environmental landmarks: sighted animals predominantly use visual signals, whereas blind animals use olfactory signals. Additionally, there appears to be a critical period soon after eye opening in which vision is required to fully refine and mature the HD system. Finally, in the absence of both visual and olfactory cues, the HD attractor network remains intact, but lacking any environmental anchors to lock onto, preferred firing directions of HD cells continuously drift over time, independent of the specific direction an animal faces.

### External (allothetic) sensory cues are required for stabilizing the head direction system

The essential input to the HD system is thought to be vestibular – neurotoxic lesions of the vestibular nerve abolished HD cell tuning in ADn (Stackman and Taube, 1997; Muir et al., 2009). In contrast, though vision is known to play a strong role in landmarking the HD cell system (Taube, 2007), since many studies have found relatively stable HD cell tuning following removal of familiar visual cues or when animals were placed in the dark (Blair and Sharp, 1996; Chen et al., 2016; Butler et al., 2017; Dannenberg et al., 2020), it has been argued that animals only require self-generated idiothetic sensory cues for stabilizing HD cell tuning (Goodridge et al., 1998; Stackman et al., 2003; Taube, 2007). Contrary to this idea, our data – in blind animals following olfactory ablation, and in sighted animals placed in the dark following olfactory ablation – reveal total loss of HD cell tuning stability in the absence of visual and olfactory cues. This means that animals require externally generated, allothetic environmental cues processed via either visual or olfactory systems, or both together, in order to stabilize the preferred firing direction of HD cells in ADn. Our results also indicate that idiothetic cues alone (for instance arising from vestibular, proprioceptive or motor efference copy systems) are insufficient to stabilize HD cell tuning in ADn. Furthermore, our results suggest that previous studies describing stable HD cell tuning in the absence of visual inputs likely arose from either insufficient removal of all visual cues or from the presence of olfactory cues. Thus, it appears that intact vestibular and visual/olfactory signals are both required for stabilizing HD cell tuning in ADn.

Aside from visual and olfactory inputs, our results rule out a contribution of vibrissal signals in stabilizing HD cell tuning, consistent with recent findings from somatosensory cortex (Long and Zhang, 2021). Next, since our experiments were done in the presence of white noise, we did not directly test possible auditory control of HD cell tuning. As such, future studies will be required to test for the possible use of auditory cues in stabilizing HD cell tuning in blind mice, though previous work in blindfolded rats suggests that auditory cues do not exert a strong effect on HD cell tuning (Goodridge et al., 1998). However, it should be noted that a recent study finds that some mice can echolocate (He et al., 2021).

### Olfaction, HD cell tuning, and cognitive maps

Our results indicate that vision and olfaction can be used, either independently or in tandem, to stabilize HD cell tuning. Visual inputs are believed to enter the HD system via inputs from visual cortex to postsubiculum and retrosplinial cortex (Taube, 2007). With respect to olfactory inputs, direct projections from the olfactory bulb and piriform cortex to the entorhinal cortex (Agster and Burwell, 2009; Chapuis et al., 2013) could in turn reach ADn via postsubiculum, which is reciprocally connected to the entorhinal cortex and ADn (van Groen and Wyss, 1990). Future studies examining these pathways in more detail in both sighted and blind mice will be required to develop a better understanding of the circuitry involved.

We find that, at least in our open field arena, vision promotes the most refined HD cell tuning, with olfaction driving slightly less refined HD tuning in blind mice. Furthermore, blind mice appear more capable of using olfaction to stabilize HD cell tuning than sighted animals: for blind mice, olfactory cues (upon floor rotation) exerted control on HD cells at levels similar to the control of visual cues in sighted animals, whereas sighted animals placed in the dark exhibited significantly less robust olfactory cue anchored rotation. Our finding of weak olfactory guided HD cell control upon floor rotation in sighted mice is consistent with a previous study in sighted rats (Goodridge et al., 1998). Overall, our results are consistent with a foraging study in rats that posited a sensory hierarchy in spatial navigation, with vision being at the top of the hierarchy, followed by olfaction (Maaswinkel and Whishaw, 1999).

Outside the HD cell system, a recent study has shown that spatial maps can be found in piriform cortex during a spatial-olfactory learning task, with spatially encoding piriform cortex cells being linked to hippocampal theta (Poo et al., 2021). Another recent study found that olfactory cues can modulate place cell responses in hippocampus and aid in path integration (Fischler-Ruiz et al., 2021). Consistent with these findings, earlier studies in rats found intact place cells in blind animals (Save et al., 1998), as well as a contribution of olfactory inputs to place cell stability (Save et al., 2000). Therefore, seeing as olfactory inputs can enable robust and stable place cell and HD cell coding in the brain, olfaction needs to be considered an important sensory system for the generation of cognitive maps.

What olfactory cues do mice use to landmark their HD system? Previous work indicates that rodents can use a variety of odors, including self-generated odors, conspecific odors, and other non-animal odors to guide spatial navigation (Wallace et al., 2002; Fischler-Ruiz et al., 2021; Poo et al., 2021). In our experiments, for each mouse, its first open field exposure was started on a new floor, which had never been explored by another mouse, meaning the mice had to use cues intrinsic to the floor or that were self-generated.

### Vision around the time of eye opening is required for refining HD cell tuning

Hubel and Wiesel discovered a critical period during development for ocular dominance plasticity in visual cortex (Wiesel and Hubel, 1963), and similar critical periods have been shown to exist for many other sensory systems and behaviors (Berardi et al., 2000). Here, we provide evidence of a critical period in the refinement and maturation of the HD system in ADn that depends on visual inputs in the period shortly after eye opening – the evidence being that rd1 mice, who have normal vision upon eye opening before going blind around P30, have more refined HD cell tuning as adults than congenitally blind Gnat1/2^mut^ mice. Both rd1 and Gnat1/2^mut^ mice go blind due to problems with retinal photoreceptors. rd1 mice go blind as a result of photoreceptor degeneration caused by a mutation in phosphodiesterase in rod photoreceptors (due to a mutation of the *Pde6B* gene), that initially leads to rod death followed by cone death, and this is a commonly used mouse model of retinitis pigmentosa (Chang et al., 2002). Gnat1/2^mut^ mice are blind as a result of mutations in both rod and cone forms of the alpha subunit of the G-protein transducin (due to mutations in both *Gnat1* and *Gnat2* genes), resulting in non-functioning rod and cone photoreceptors (see Methods). As the mutated genes in both mouse lines are predominantly expressed in photoreceptors, it is likely that the difference in HD cell tuning in blind adult Gnat1/2^mut^ vs. rd1 mice is a direct result of the timing of the onset of vision loss (congenital vs. ∼P30). We thus propose that vision around the time of eye opening allows the HD cell system to stabilize (i.e. upon eye opening, vision enables HD cells to exhibit heightened stability in their preferred direction tuning (Bjerknes et al., 2015; Tan et al., 2015)) and this stability results in refinement and maturation of the HD network, such that though both rd1 and Gnat1/2^mut^ are equally blind as adults, rd1 mice have significantly more refined HD cell tuning. These findings are consistent with multisensory studies in superior colliculus which showed the importance of vision during development for enabling the generation of normal auditory space maps (King et al., 1988; King and Carlile, 1993; Wallace and Stein, 2007).

### Attractor dynamics in the presence and absence of anchoring external sensory inputs

The HD network is often modeled as a continuous ring attractor (Skaggs et al., 1995; Peyrache et al., 2015; Chaudhuri et al., 2019; Angelaki and Laurens, 2020). Evidence in favor of an attractor network is plentiful, from the finding that the relative difference in preferred directions between a given pair of HD cells is maintained across different environments and during visual cue rotation experiments (Taube et al., 1990b), to the finding that HD cells that fire coherently during spatial navigation continue to do so during sleep (Peyrache et al., 2015), to the finding that before eye opening – when HD cell tuning curves are broad – pairs of cells exhibit similar coherence during spatial navigation as is seen in the adult HD system (Bjerknes et al., 2015; Tan et al., 2015; Bassett et al., 2018). Furthermore, a recent manifold analysis has validated a one-dimensional ring attractor as a robust description of the rodent HD network (Chaudhuri et al., 2019). Our results in blind mice are consistent with those of a continuous ring attractor. Performing a manifold analysis revealed a one-dimensional ring attractor similar to that found in normally sighted animals. Additionally, upon ablating OSNs in blind animals, while this completely removed stability of HD cell preferred direction, attractor dynamics remained intact.

We find that – for normally sighted and blind animals – the removal of both visual and olfactory inputs causes the ‘hill of activity’ within HD attractor network to drift independently from the animal’s true heading direction (i.e. the preferred direction of all simultaneously recorded cells drifts coherently). For a given animal, over a 10-minute recording session, the drift could be predominantly in either CW or CCW direction, or exhibit no strong overall directional bias. For blind animals, there appeared to be a bias towards CW drift. For some animals, there was a significant correlation between angular head velocity and angular drift velocity of PFDs.

A similar drift in HD cell preferred firing direction appears to have been described in a previous study recording from HD cells in the lateral dorsal nucleus in rats placed in the dark, though those animals were placed in a radial maze, making it difficult to draw exact parallels to our results (Mizumori and Williams, 1993). In a more recent study in mice placed in the dark, it was reported that approximately 40% of HD cells became unstable in the dark, with one example HD cell recording being shown where the instability resulted in the preferred firing direction exhibiting a drift similar to what we describe (Yoder and Taube, 2009). However, a follow-up paper from the same lab showed no effect of dark exposure on HD cell stability, and instead found that optogenetic silencing of the nucleus prepositus hypoglossi (NHP) – which relays vestibular signals to the HD system – was required in tandem with dark exposure to cause the preferred direction of a subset of HD cells recorded in ADn to drift over time (Butler et al., 2017). This latter study also found that preferred firing direction drift could occur in either CW or CCW directions, and was sometimes correlated with the animal’s head turns (Butler et al., 2017). Such drift in HD preferred direction may be related to the effect described for HD cells prior to eye opening, where the broadness of tuning curves appeared to result from an under-signalling of angular head velocity (Bassett et al., 2018), though the drift we report in the absence of both visual and olfactory inputs is significantly stronger, and in many animals we found no clear relationship between preferred direction drift and angular head velocity. Thus, future work is required to understand the circuit and synaptic basis whereby different sensory inputs enable tuning within the HD attractor network to stabilize, and how perturbations to different sensory modalities can alter the input to the HD network and result in drift.

### Relating head direction cell recordings in blind rodents to spatial cognition in visually-impaired humans

While vision is often considered the most important sensory system for guiding spatial cognition in humans, numerous studies have shown that visually impaired individuals maintain robust spatial cognition (Loomis et al., 1993; Thinus-Blanc and Gaunet, 1997; Tinti et al., 2006; Schinazi et al., 2016). Unlike in blind mice – where olfaction appears to be the most important sensory system for guiding spatial awareness following vision loss – in visually-impaired humans the auditory system is generally thought to be a more important contributor to vision-independent spatial awareness, with blind humans appearing to exhibit enhanced sound localization compared to sighted subjects (Lessard et al., 1998) and performing echolocation via self-produced sounds (Kellogg, 1962; Stroffregen and Pittenger, 1995; Milne et al., 2014). However, there is emerging evidence that humans can also effectively use olfaction to inform spatial cognition (Porter et al., 2007; Jacobs et al., 2015; Hamburger and Knauff, 2019), meaning that olfaction could be used following vision loss to aid in spatial awareness and navigation (Koutsoklenis and Papadopoulos, 2011). Next, while there is active debate surrounding the extent to which visual experience in humans is required for developing normal spatial cognitive abilities (Pasqualotto and Proulx, 2012; Schinazi et al., 2016), some studies have indicated that late-blind individuals perform better on certain spatial cognition tasks than congenitally blind individuals (Rieser et al., 1980; Herman et al., 1983; Rieser et al., 1986; Pasqualotto and Newell, 2007), reminiscent of our findings regarding HD cell tuning being more refined in ‘late-blind’ rd1 mice compared to congenitally blind Gnat1/2^mut^ mice. Thus, our findings related to HD cells in sighted and blind mice are likely highly relevant for understanding the neural underpinnings of spatial cognition in humans following vision loss.

## Methods

### Animals

All procedures were performed in accordance with the Canadian Council on Animal Care and approved by the Montreal Neurological Institute’s Animal Care Committee. Three strains of mice were used in this study: wild type mice (C57Bl/6; Charles River strain code 027), rd1 mice (also known as C3H; The Jackson Laboratory #000661), and Gnat1/2^mut^ mice (Gnat2^cpfl3^ Gnat1^irdr^/Boc mice; The Jackson Laboratory #033163). All mice were adults (2-4 months old; P75-125) weighing between 20 to 32 grams. Mice of both sexes were included.

### Surgery and implantation

Mice were anesthetized with a cocktail containing fentanyl (0.05mg/kg), medetomidine (0.5mg/kg) and midazolam (5mg/kg) (Hillier et al., 2017), and a craniotomy was performed above ADn for probe implantation. A conductive wire was inserted into the cerebellum to serve as a reference. After attaching the reference wire, a 4-shank silicon probe (Neuronexus Inc. Ann Arbor, MI; 200 µm inter-shank spacing) with 8 recording sites on each shank (probe model Buz32) was lowered into the brain towards ADn based on the following stereotactic coordinates: antero-posterior (AP) −0.4 mm; medio-lateral (ML) −0.76 mm; dorso-ventral (DV) 2.16 mm. The base of a movable drive holding the silicon probe was then fastened to the skull using dental acrylic cement and a light-cure adhesive (Kerr OpitBond Universal Unidose) to allow for stable recordings during awake open field sessions.

### Electrophysiological recordings

After recovery (5-7 days post-surgery), during sleep the probe was advanced daily in steps of < 300 µm in the home cage until HD units were detected in ADn. Prior to open field recording sessions, screenings were done during sleep in the homecage. Preliminary detection of putative HD cells was based on inspection of auto-correlograms of recorded units, as previously described (Viejo and Peyrache, 2020). Once putative HD units were detected, experiments were conducted by placing animals in an open field dark cylindrical arena (60 cm in diameter) with a single white visual cue placed on the wall above the animal’s reach. Open-field recordings lasted for 10 minutes per session. The neurophysiological signals were acquired at 20 kHz using an Intan RHD2000 Recording System (16-bit, analog plexin). The raw neuronal signal was high-pass filtered and processed with an automated spike sorting algorithm to extract single units (Kilosort2; Pachitariu et al., 2016). Isolated units were manually curated in Klusters (Hazan et al., 2006) based on auto-correlograms and spike waveforms. For Figure 4E where cells were recorded over a 2-day period, the sessions were merged to form a single continuous recording before spike sorting to ensure that the same units were maintained for further analysis.

### Position tracking

For position tracking, an infrared-based (850 nm IR) camera recording system equipped with 8 cameras recording at 120 FPS were mounted above the recording arena to capture the movements of the animal (Optitrack Flex 13). Four reflective infrared markers were attached to the animal’s head-stage for tracking (6.4mm Optitrack M3 Markers). After recording, Motive motion capture software (Optitrack) was used to extract both the direction and position of the head relative to the environment in 3 dimensions. It should be noted that for dark experiments, we found that sighted mice could see the visual cue in ‘the dark’ when we used the 850 nm IR illumination that came with the Optitrack cameras (i.e. HD cells followed the visual cue in ‘the dark’ when it was moved to a new location (2 animals, data not shown)). This is consistent with findings from a recent paper about red light perception in rats (Nikbakht and Diamond, 2021). As such, for dark experiments, we installed a custom-made array of 940nm infrared lights (Adafruit, Product # 388) that we used for illumination and marker detection. Importantly, under 940 nm IR illumination HD cells did not exhibit visual cue controlled changes in their preferred direction.

### Standard HD cell classification

To identify HD cells, as mentioned above, we first used auto-correlograms to localize the electrode in ADn and identify putative HD cells located ADn (Viejo and Peyrache, 2020). Subsequently, for all cells recorded in ADn, tuning curves were constructed by aligning spike times with the closest head direction point in the horizontal plane. Spikes were then counted for bins of 6°. To correct for angular sampling bias, spike rates were normalized by the time spent in each angular bin, as previously described (Peyrache et al., 2015). Units that passed a Rayleigh test for significant non-uniform circular distribution (p <0.0001), had a z-test score > 50, and a peak rate > 1Hz, were classified as HD cells.

### Gradient boosted trees (XGBoost) model for HD cell classification

Owing to the loss of stable HD tuning in blind animals following olfactory sensory neuron ablation, cells recorded in ADn in this condition were classified as HD/non-HD cells using the XGBoost model (Chen & Guestrin, 2016), implemented in Python. First, in control animals (sighted in the light (WT_L_) and blind) we used the standard HD cell classification method (see above) to define cells in ADn as either HD or non-HD cells. Next, we generated auto-correlograms from these HD and non-HD cells using 2ms bins. Since auto-correlograms are symmetrical, for further processing we only used the half corresponding to the positive lag. Using default parameters, the XGBoost model was trained on a subset (80%) of the pre-defined (i.e. labelled as HD or non-HD cells) auto-correlograms with a 10-fold validation, and then the performance was evaluated on the held-out test data (20%). The model performance on the test data was 85.3%. Next, the trained model was used to classify ADn cells recorded in blind animals following OSN ablation as either HD or non-HD cells based on their auto-correlograms.

### Visual cue rotation

The extent to which the single white cue-card on the arena wall controlled the preferred firing direction (PFD) of HD cells was tested by rotating the visual cue by either 90° or 180°. Following a standard session where the baseline PFDs for all simultaneously recorded HD cells was established, the animal was removed from the arena and disoriented, as previously described (Taube, 1995). The animal was then reintroduced to the arena after the visual cue was rotated. For all visual cue rotation experiments, the floor of the arena was thoroughly cleaned with 70 % ethanol to eliminate olfactory cues on the floor before the animal was reintroduced. The visual cue control measure (gain) shown in Figure 1G was computed as follows:

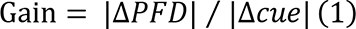

where, Δ*PFD* is defined as the change in the preferred firing direction (in degrees) for each HD cell following reintroduction into the area, and Δ*cue is* defined as the extent (either 90° or 180°) that the visual cue was rotated.

### Floor rotation

To assess floor controlled changes in PFDs of HD cells, a protocol similar to visual cue rotation described above was adopted, except where the floor of the arena was rotated. For these experiments, to avoid possible interference of odor markings across different animals, a brand new floor was used for each mouse. At the same time, to give mice the opportunity to use olfactory odors on the floor, the floor of the arena was not cleaned between exposures. The extent to which floor rotations influenced the PFD of HD cells across animals was computed using equation (1).

### Olfactory sensory neuron (OSN) ablation

To eliminate (or at least greatly reduce) the sense of smell, OSNs were chemically ablated based on established methods (McBride, 2003; Norwood et al., 2019). In brief, mice were anesthetized with isoflurane after which a blunted 33-gauge needle was used to administer 20μl of ZnSO4 (10 % in sterile H_2_O) in both nostrils. After intranasally administering the chemical, mice were inverted to drain the excess fluid from the nasal cavity. Behavioral and electrophysiological recordings were carried out at least 24hrs following OSN ablation.

### 2-Chamber olfactory test

To test the efficacy of OSN ablation, in some treated animals we tested their sense of smell using a 2-chamber odor test. Animals were first habituated to the chambers for 5 minutes each day over a period of 3 days, with no odors added to the chambers. After habituation, testing began by placing two filter papers infused with 10µl of 3-methyl-1-butanethiol (aversive odor; Sievert and Laska, 2016) in a randomly selected chamber, and two filter papers infused with 10µl of distilled water (neutral odor) in the other chamber. All mice were placed in the neutral chamber at the start of the test and were monitored for 10 minutes using an over-head camera to record their baseline occupancies in both chambers. Post-OSN ablation, the test was repeated, and compared to the baseline. The chamber preference (CP) score in Figure 4 was computed as follows:

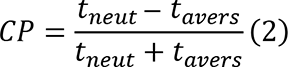

where, *t_neut_* and *t_avers_* refers to the time spent in the neutral and aversive side respectively. Chamber preference scores of −1 and 1 represent strong preference for the aversive and neutral side respectively, where the animal spent the entire time in one chamber.

### Drift analysis

Following the loss of vision and olfaction, PFD drift within the HD population was computed as follows: First, the PFD of simultaneously recorded HD cells was computed each time the animal completed a full head turn. The resulting change in PFDs over the 10-minute recording session was averaged across cells and filtered with a Gaussian kernel (s.d.=1.5) to provide an estimate of the drift over time.

### Drift direction index (DDI)

To quantify the consistency in drift direction, we computed a drift direction index by taking the difference of CW and CCW drift counts normalized to the total number of drift events in each session (equal to the number of 360° head turns in a 10 minute session). The DDI value for each session thus ranges from −1 to 1, where −1 and 1 represent consistent drifts in the CCW and CW direction respectively.

### Bayesian decoding

To predict the HD of the animal given the spiking activity of HD cells within a given time window, a Bayesian decoding algorithm (Zhang et al., 1998) was used to compute the posterior probability distribution using the formula:

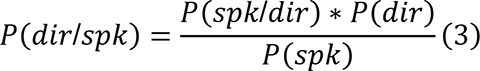

where, *dir* represents the set of possible angular head directions and *spk* represents the spike counts of simultaneously recorded HD cells within a 200ms time window in this instance. Bayesian decoding was used to predict the HD of animals in the second half of a recording session based on priors generated from tuning curves in the first half. Mean absolute decoding error was computed as the absolute difference between the decoded angle and the actual angle over the decoded time window.

### ISOMAP projection

Isomap analysis was performed on sessions with at least 12 HD cells owing to the large number of data points needed for the unsupervised algorithm to work efficiently (Xiang and Gong, 2005). Similar to Viejo and Peyrache (2020), a square root transformed 200ms binned-spikes from all simultaneously recorded units in ADn served as inputs to the ISOMAP algorithm (Tenenbaum et al., 2000), implemented in python (Pedregosa et al., 2011). The number of neighbors was set to 50. The resulting output of the algorithm formed the basis of the low dimensional ring manifolds shown in Figure 6 and Supplementary Figure 6. The color code of the population vector on the ring manifold was mapped onto the corresponding actual HD for each corresponding time bin.

### Quantification of ADn units

To ensure that cells counted on recording shanks were recorded in ADn, the following was implemented: first, auto-correlograms were generated from all cells on shanks that picked at least 1 HD cell. Second, a Fourier transform was applied to the auto-correlograms to identify and exclude theta (4-8Hz) modulated cells. Although some studies have reported the presence of theta modulation in surrounding anterior thalamic structures (Jankowski et al., 2013; Tsanov et al., 2011), a recent study shows that cells in ADn are not theta modulated (Viejo and Peyrache, 2020). Hence the total counts of ADn cells were derived from all non-theta modulated units. Finally, the proportion of ADn units characterised as HD cells was derived from the total number of units classified as HD cells (see Methods: HD classification) divided by the total number of cells recorded in ADn. The difference between proportions was tested using a 2-sample Z-test for proportions as previously described (Bassett et al., 2018).

### Analysis of head direction cell

Mean firing rate: The temporal average of spike counts. Peak firing rate: The normalized maximum spike counts in a second. Mean vector length: The circular spread of spikes, with 0 and 1 representing strong uniform and non-uniform circular spread respectively. In Figure 6D and Supplementary Figure 6D, for each cell, the mean vector length for the short intervals was computed by averaging the vector length values across all 360° head turns in a session. Tuning width: The full width at half max of a cell’s tuning curve. Mutual information: An estimate of the directional information relating the firing rate of a cell to a given direction. The formula (Skaggs et al., 1996) is given as follows:

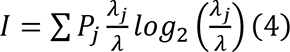

where, *I* refers to the information content in bits/spike*, P_j_* is the probability of occupying bin *j, λ_j_* is the mean firing rate in bin *j,* and *λ* is the in-session mean firing rate of the cell. Stability: The Pearson correlations of the tuning curve of cells from the 1^st^ and 2^nd^ half of a 10-minute recording session. In Figure 1D, stability was defined as the Pearson correlations of the mean PFDs of simultaneously recorded cells in the 1^st^ and 2^nd^ half of a recording session pooled across animals. Similarly, in Figure 1E, stability was assessed based on the Pearson correlations of mean PFDs of exposure 1 and 2 across all recorded animals.

### Statistical analyses

The statistical test used for every specific comparison is directly stated in the text or figure legend. For hybrid violin/box-plots, the violin plot shows the density distribution of all data points, whereas the box-plot shows the median, 25/75% distribution, and 5/95% distribution. For statistical comparisons, * p < 0.05, ** p < 0.01.

---

**Supplementary Figure 1.**
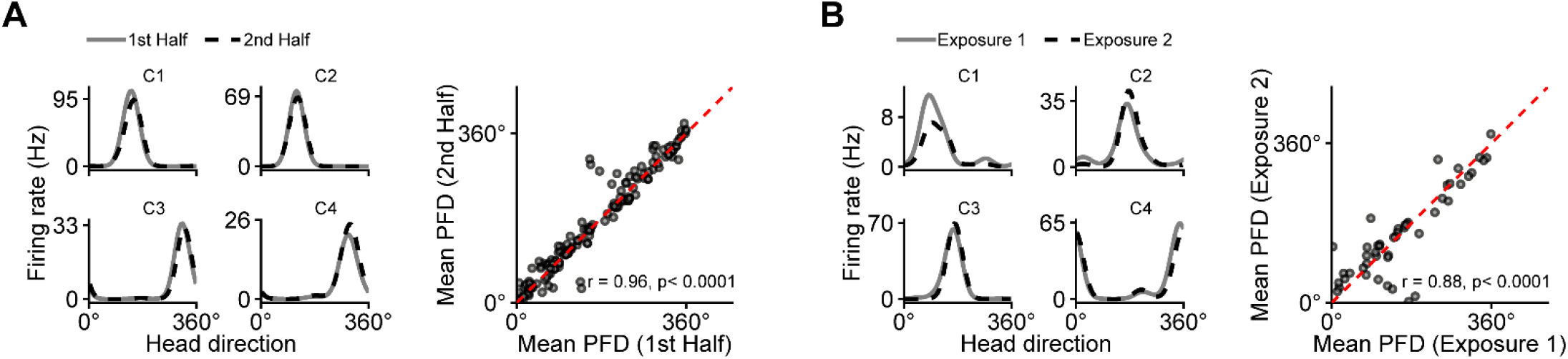
**A**, *left*, The spike rate vs. heading direction of 4 simultaneously recorded HD cells (C1-C4) in a sighted mouse during the first and last 5 minutes of a 10-minute recording session. **A**, *right*, For all HD cells (128 HD cells across 9 animals), the mean PFD is compared between the first and last 5 minutes. The pre-post tuning similarity was tested with a Pearson correlation, and the resulting *r* value is shown. **B**, The same as in **A**, except comparing HD cell tuning across successive 10-minute exposures to the same open field arena (48 HD cells across 4 animals).

**Supplementary Figure 2.**
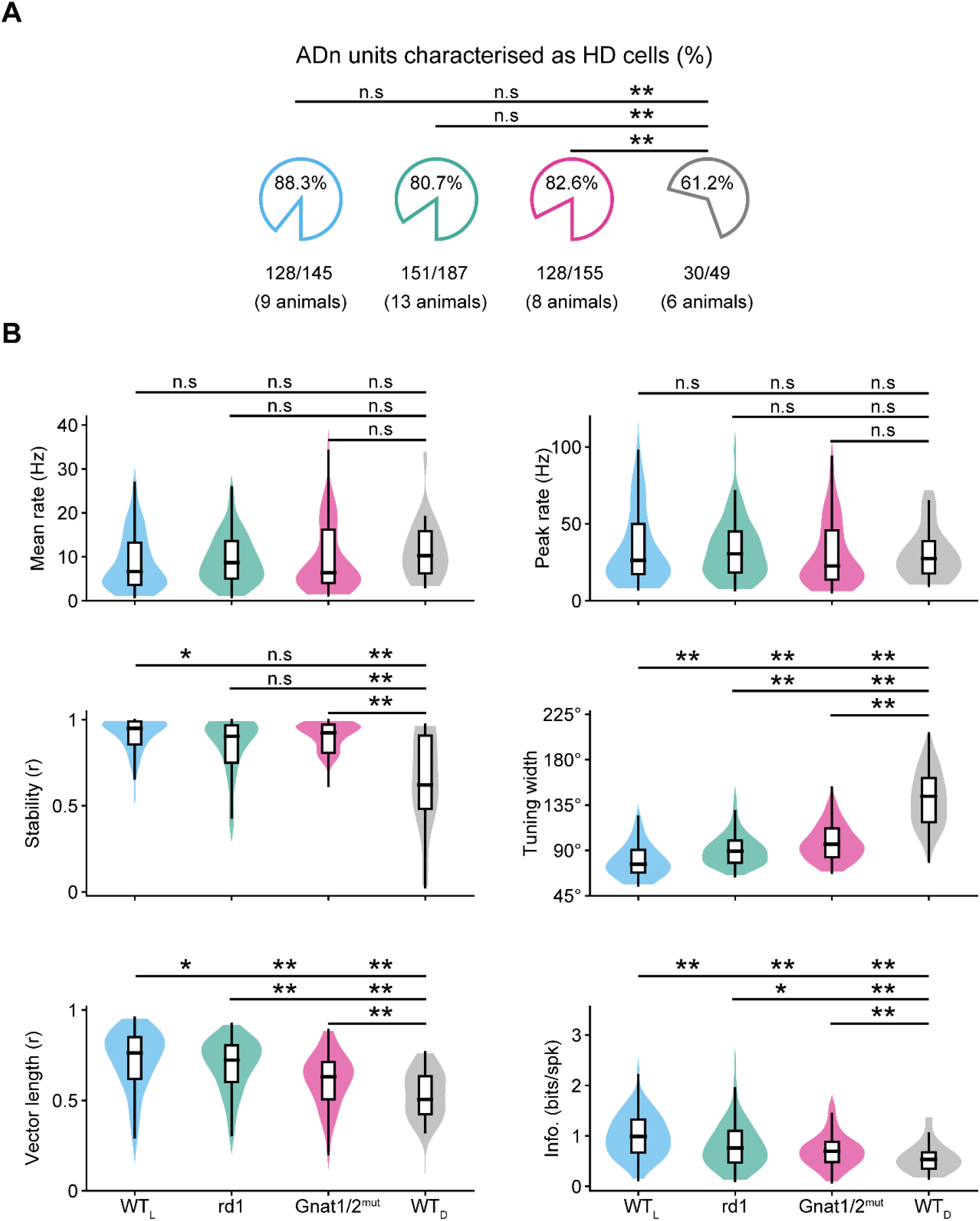
Replots from Figure 2 and 3, for direct comparison between all different mouse lines and manipulations in Figures 2 and 3. **A**, For the 4 different groups of mice – wildtype in light (WT_L_; blue), rd1 (blind by ∼ P30; green), Gnat1/2^mut^ (congenitally blind; pink) and wildtype in the dark (WT_D_; grey), the average percent of cells recorded in ADn that passed the criterion to be designated as HD cells (see Methods) are compared. Statistical differences were calculated using the Z-test for Proportions. **B**, For the 4 different groups of mice in **A**, several metrics of HD cell responses are compared. For each graph, data are shown as hybrid violin/box plots (see Methods). Statistical differences were calculated using the Kolmagorov-Smirnov Test with Bonferroni Correction for multiple comparisons.

**Supplementary Figure 3.**
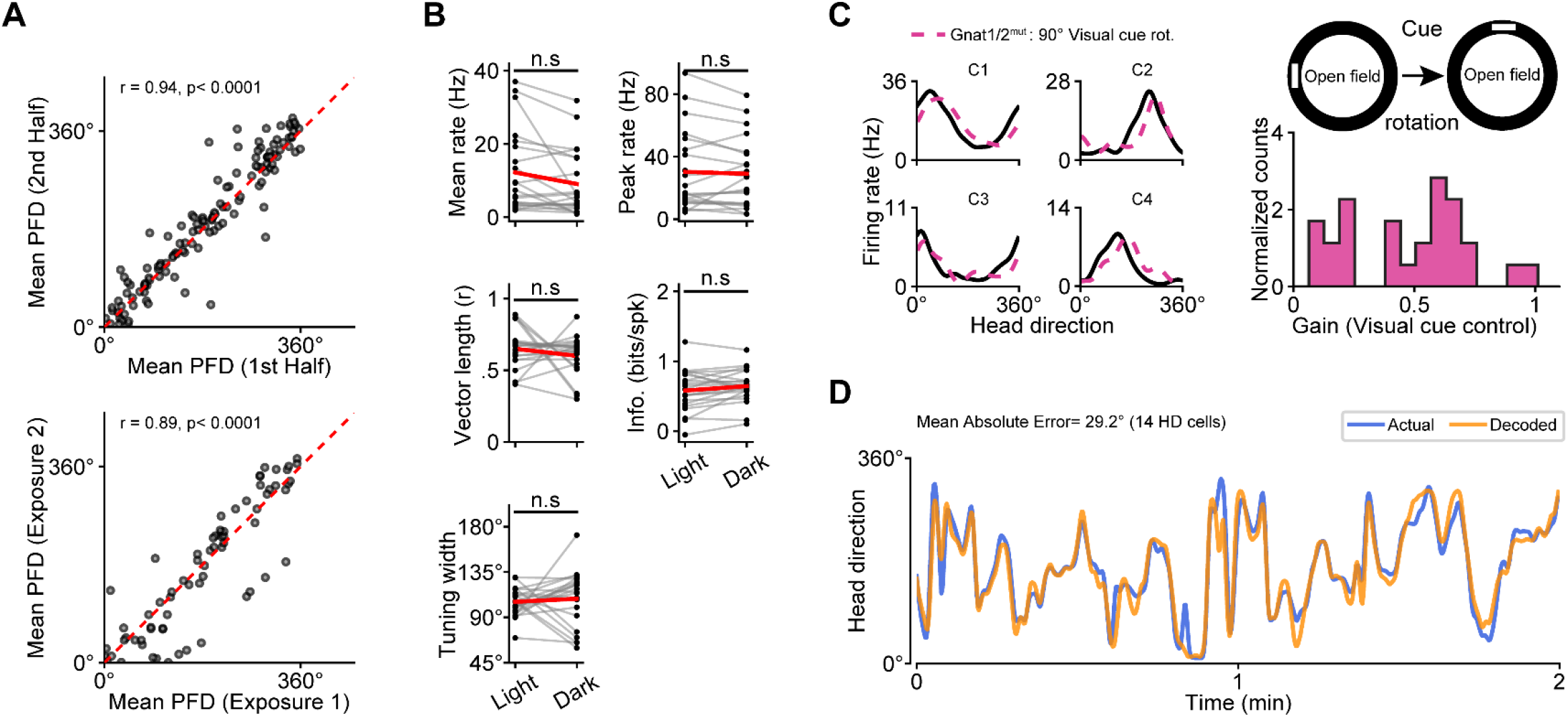
**A**, *top*, For all HD cells from Gnat1/2^mut^ animals (128 HD cells across 8 animals), the mean PFD is compared between the first and last 5 minutes. The pre-post tuning similarity was tested with a Pearson correlation, and the resulting *r* value is shown. **A**, *bottom*, The same as in **A**, except comparing HD cell tuning across successive 10-minute exposures to the same open field arena (70 HD cells across 5 animals). **B**, Several metrics characterizing HD cells in Gnat1/2^mut^ mice are compared during light vs. dark exposure (21 HD cells across 3 animals). Statistical differences were calculated using the Wilcoxon Signed-Rank Test. n.s = not statistically different. **C**, *left*, The spike rate vs. heading direction of 4 simultaneously recorded HD cells in a Gnat1/2^mut^ following a 90° visual cue rotation (dotted line). **C**, *right, A* schematic indicating the nature of visual cue rotation experiments, and a histogram showing the extent of control the visual cue rotation exerted on the PFD of HD cells in Gnat1/2^mut^ mice (28 HD cells across 5 animals). Note that for visual cue rotation experiments, we cleaned the floor in between exposures (i.e. the mouse was removed from the room, the cue was moved, the floor was cleaned, and then the animal was placed back in the arena – and this is why the stability is not as robust in panel **A**, *bottom*). **D**, Analysis comparing the actual heading direction of a Gnat1/2^mut^ mouse over time (blue) to the heading direction predicted by a Bayesian decoder (orange, see Methods).

**Supplementary Figure 4.**
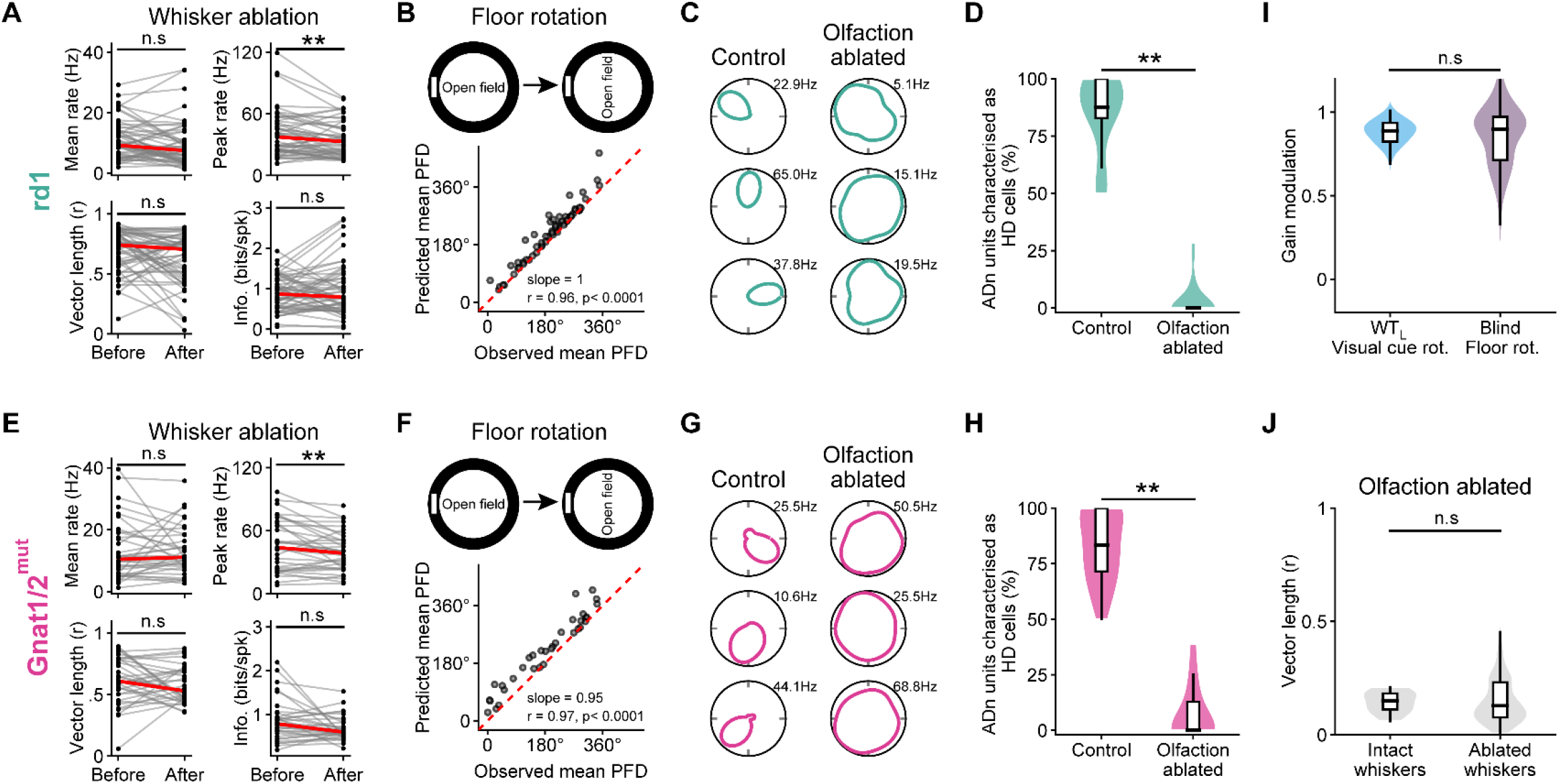
**A-H** Similar to Figure 4A, except separate analyses are shown for rd1 and Gnat1/2^mut^ mice. **I**, The extent to which visual cue/floor rotation exerted control on the PFD of HD cells (visual cue for sighted animals in the light (WT_L_) and floor rotation for blind animals (pooled rd1 and Gnat1/2^mut^ mice)), measured as the gain between the extent of HD cell PFD rotation compared to the extent that the visual cue/floor was rotated. **J**, Comparison of the vector length of HD cells recorded in blind mice following olfaction ablation, with and without whiskers. Statistical difference was tested with the Mann-Whitney U Test. Intact whiskers = 29 HD cells; Ablated whiskers = 77 HD cells.

**Supplementary Figure 5.**
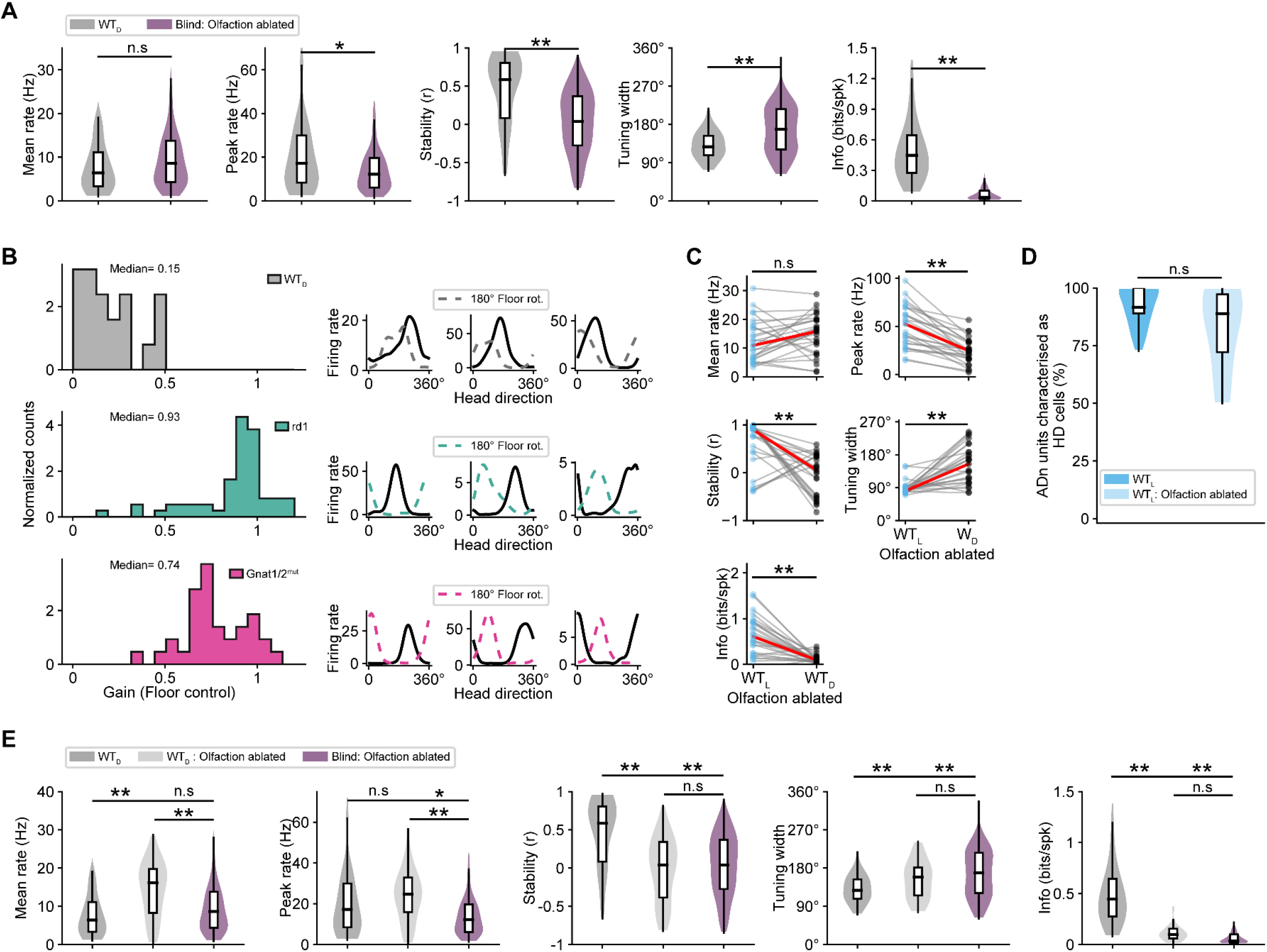
**A**, Related to Figure 5A, additional comparisons of HD cell metrics between wild type in the dark (WT_D_) and olfaction ablated blind mice. **B**, *left*, Histogram showing the extent of control the floor rotation (gain) exerted on the PFD of HD cells for each group: sighted animals in the dark (WT_D_), rd1 and Gnat1/2^mut^ mice. WT_D_ = 20 HD cells across 4 animals, rd1 = 60 HD cells across 5 animals, Gnat1/2^mut^ = 34 HD cells across 4 animals. **B**, *right*, For each of the different groups of mice listed on the left (see color legend), the spike rate vs. heading direction of 3 example HD cells in control condition (solid black line) and following a 180° visual cue rotation (dotted line). **C**, Several metrics of HD responses from sighted animals placed in both light vs. dark environments following olfactory sensory neuron ablation. n = 27 HD cells across 5 animals. Statistical differences were calculated using the Wilcoxon Signed-Rank Test. n.s = not statistically different. **D**, Comparison of the % of ADn cells characterized as HD cells in sighted animals placed in light before and after olfactory sensory neuron ablation. Statistical difference was tested using the Z-test for Proportions. **E**, Comparison of HD cell metrics between sighted animals placed in the dark prior to OSN ablation (WT_D_), post-OSN ablation (WT_D_: Olfaction ablated) and olfaction ablated blind mice (Blind: Olfaction ablated). Statistical differences were calculated using the Kolmagorov-Smirnov Test with Bonferroni Correction for multiple comparisons. WT_D_= 42 HD cells across 6 animals; WT_D_: Olfaction ablated= 32 HD cells across 5 animals; Blind: Olfaction ablated= 106 HD cells across 9 animals.

**Supplementary Figure 6.**
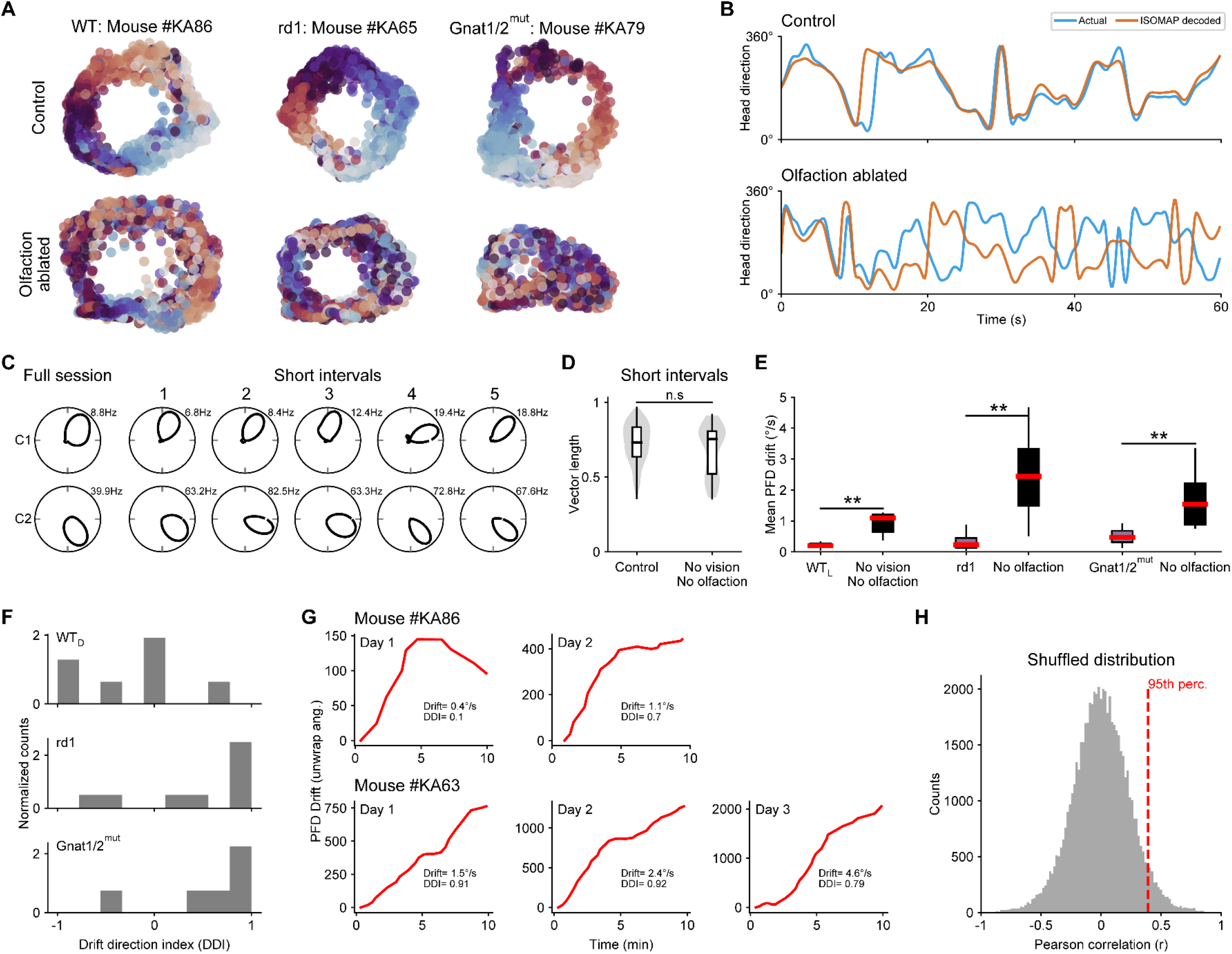
**A**, *top*, Example Isomap plots outlining the 1-dimensional ring manifold of the HD cell network in sighted animals in the light (WT_L_, *left*), rd1 mice (*middle*) and Gnat1/2^mut^ mice (*right*). **A**, *bottom*, The same as above, except following olfactory ablation (also note that the WT example is now from an animal placed in the dark (WT_D_)). **B**, For an example rd1 mouse, ‘decoding’ is shown using the Isomap ring projection, with the animal’s actual heading direction being compared to the ring-decoded head direction (see Methods) in control (*top*) and following olfactory ablation (*bottom*). **C**, Example polar plots for 2 simultaneously recorded HD cells in a sighted mouse in the light, either calculated over the entire 10-minute recording session (*left*) or over shorter timescales (right) with each successive epoch computed upon successive 360° head turns (for comparison to Figure 6C). **D**, Comparison between the responses of HD cells computed on short timescales (each time the animal makes a 360° head rotation) compared between control animals (combined sighted wildtypes in the light and blind mice; n = 407 HD cells) and animals with No vision and No olfaction (n = 133 HD cells). Statistical difference was tested with the Mann-Whitney U Test. **E**, The average velocity of drift in the HD population is compared in different groups of control mice, and the same mice without both vision and olfaction. Statistical difference was tested with the Mann-Whitney U Test. WT_L_ (9 animals) vs WT_D_: Olfaction ablated (5 animals); rd1 (13 animals) vs rd1: Olfaction ablated (5 animals); Gnat1/2^mut^ (8 animals) vs Gnat1/2^mut^: Olfaction ablated (4 animals). **F**, Histogram showing the extent that the overall drift direction was either CW (positive values) or CCW (negative values) for different groups of mice with no vision and olfaction (see Methods for drift direction index description). WT_D_= 5 animals; rd1=5 animals; Gnat1/2^mut^= 4 animals. **G**, For 2 different animals without vision and olfaction, the HD cell population drift (i.e. the average PFD drift for all simultaneously recorded HD cells) is shown across multiple days. **H**, Histogram of shuffled correlations generated by shuffling AHV 10,000 times and correlating it with ADV after each shuffle (see Methods). The absolute correlation values of sessions indicated in red (see Figure 6J) exceeded the 95^th^ percentile of the null distribution (r value = 0.39).

## Acknowledgements

We thank Drs. R. Tong and G. Viejo for critical discussions on this project. We thank A. Villemain for maintaining mouse colonies. We acknowledge the following funding sources: Jean Timmins Costello fellowship and Healthy Brains for Healthy Lives fellowship to K.A.; Canada Research Chairs to A.P and ST.; Alfred P. Sloan Foundation Research Fellowship, Vision Health Research Network Pilot Project for Early-Career Investigators Grant and a Canadian Institutes of Health Research Project Grant to S.T.

## Contributions

Experiments were designed by K.A., A.P. and S.T. Experiments and analyses were performed by K.A. Figures were generated by K.A. The paper was written by K.A and S.T.

## Data and code availability

All data and code are freely available upon request.

